# Fast variance component analysis using large-scale ancestral recombination graphs

**DOI:** 10.1101/2024.08.31.610262

**Authors:** Jiazheng Zhu, Georgios Kalantzis, Ali Pazokitoroudi, Árni Freyr Gunnarsson, Hrushikesh Loya, Han Chen, Sriram Sankararaman, Pier Francesco Palamara

**Affiliations:** Department of Statistics, University of Oxford, Oxford, OX1 3LB, UK; Wellcome Sanger Institute, Wellcome Genome Campus, Hinxton, CB10 1SA, UK; Department of Epidemiology, Harvard T.H. Chan School of Public Health, Boston, 02115, Massachusetts, USA; Centre for Human Genetics, University of Oxford, Oxford, OX3 7BN, UK; Human Genetics Center, Department of Epidemiology, Human Genetics and Environmental Sciences, School of Public Health, The University of Texas Health Science Center at Houston, Houston, 77030, Texas, USA; Center for Precision Health, McWilliams School of Biomedical Informatics, The University of Texas Health Science Center at Houston, Houston, 77030, Texas, USA; Department of Computer Science, UCLA, Los Angeles, 90095, California, USA; Department of Human Genetics, David Geffen School of Medicine, UCLA, Los Angeles, 90095, California, USA; Department of Computational Medicine, David Geffen School of Medicine, UCLA, Los Angeles, 90095, California, USA

**Author notes:** Contributed equally (Jiazheng Zhu, Georgios Kalantzis).

## Abstract

Recent algorithmic advancements have enabled the inference of genome-wide ancestral recombination graphs (ARGs) from genomic data in large cohorts. These inferred ARGs provide a detailed representation of genealogical relatedness along the genome and have been shown to complement genotype imputation in complex trait analyses by capturing the effects of unobserved genomic variants. An inferred ARG can be used to construct a genetic relatedness matrix, which can be leveraged within a linear mixed model for the analysis of complex traits. However, these analyses are computationally infeasible for large datasets. We introduce a computationally efficient approach, called ARG-RHE, to estimate narrow-sense heritability and perform region-based association testing using an ARG. ARG-RHE leverages a method for computing genotype-matrix products from genealogical data in sublinear time, along with scalable randomized algorithms. This enables fast estimation of variance components and their statistical significance, supports parallel analysis of multiple quantitative traits, and facilitates other linear mixed-model analyses. We conduct extensive simulations to verify the computational efficiency, statistical power, and robustness of this approach. We then apply it to detect associations between 21,159 genes and 52 blood-related traits, using an ARG inferred from genotype data of 337,464 individuals from the UK Biobank. In these analyses, combining ARG-based and imputation-based testing yields 8% more gene-trait associations than using imputation alone, suggesting that inferred genome-wide genealogies may effectively complement genotype imputation in the analysis of complex traits.

## 1. Introduction

Modern biobank initiatives have led to the collection of growing volumes of genomic and phenotypic information [54, 10, 82, 26, 38]. These datasets have been utilized in various applications, including the detection of phenotypic associations [88, 1], the study of complex trait architectures [86, 15], and the development of polygenic scores [31]. A key step in these analyses is to study the patterns of genetic and genealogical relationships among the analyzed samples, which provide insights into phenotypic and environmental variation. A wide range of approaches has been developed to analyze details of these relationships, ranging from methods that use genomic data to detect shared recent ancestry [22, 8, 60, 55, 71, 93] to approaches that reconstruct genome-wide genealogical relationships using an ancestral recombination graph (ARG) [67, 74, 30, 90].

Inferred genome-wide genealogies have predominantly been used to facilitate analyses in population genetics [67, 74, 30, 81], but have also recently been used for the study of complex traits [90, 69, 40]. One approach is to use an ARG inferred from a subset of common variants to detect the presence of unobserved genomic variation, which can then be tested for association with a phenotype of interest [52, 90]. The unobserved variation inferred from the ARG can also be used to construct a genetic relatedness matrix (GRM), which in turn can be used to estimate heritability, perform polygenic prediction, or detect associations within the framework of linear mixed models (LMMs). In this work, we adopt the terminology of [90], where ARG-derived GRMs were introduced for LMM-based complex trait analyses and referred to as ARG-GRMs. A contemporary approach [14] independently described a similar model for population genetic analyses, referred to as eGRM. Both models build on related work on the genealogical interpretation of principal component analysis [51].

When used to estimate heritability, ARG-GRMs can potentially improve the estimates by including the contribution of variants that would otherwise not be observed [90]. This strategy can also be utilized for variance component association testing [40] by constructing the GRM for a specific region of interest and assessing whether heritability significantly deviates from zero. By reducing the number of tests performed and pooling potentially heterogeneous effects within a region, as is often done in the analysis of rare variants [56, 83, 39, 63], this ARG-based variance component testing approach may lead to increased association power. However, estimating global or local heritability using the approaches of [90] and [40] requires generating and manipulating ARG-GRMs that contain *N* × *N* entries for *N* studied individuals, which becomes computationally infeasible in datasets comprising more than a few thousand individuals.

Here, we develop an approach, called ARG-RHE, that uses genealogical data and graph-based algorithms to test for association within a genomic region. ARG-RHE is highly scalable, making it applicable to ARGs inferred from large biobank datasets, and allows modeling the phenotypic effects of variants that are not genotyped or accurately imputed using sequencing reference panels. It combines a scalable ARG traversal algorithm with two existing methodological frameworks: randomized algorithms for moment-based estimation of narrow-sense heritability (RHE-mc [84, 62]) and moment-based approximations for computing tail probabilities in significance testing (FastSKAT [83, 46]). We show that matrix multiplication using the ARG scales sublinearly with sample size and enables a range of downstream applications, and we evaluate the scalability, calibration, and statistical power of ARG-RHE using simulated data. We then apply ARG-RHE to detect associations between 52 quantitative blood-related traits and an ARG inferred from SNP array data for 337,464 individuals in the UK Biobank (UKBB; [10]). We find that the associations detected using ARG-RHE are complementary to those identified by testing ∼54 million variants imputed from ∼65, 000 sequenced haplo-types in the Haplotype Reference Consortium (HRC [50]) panel, and are enriched for rare variant associations detected using exome sequencing data [28].

## 2. Methods

In this section, we first introduce the trait model and notation used throughout the work. We then describe the Randomized Haseman–Elston regression (RHE) algorithm, on which ARG-RHE is built, along with its extension for parallel analysis of multiple traits. Next, we present an algorithm for fast multiplication between the genotype matrix and an arbitrary matrix, explain how it is used to implement ARG-RHE, and discuss applications in additional domains. Finally, we describe the experimental setup used to benchmark these methods on simulated and real datasets.

### 2.1. Trait model

We assume a linear relationship between the phenotype and the genetic and environmental components,

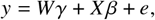

where *Y* is the phenotype vector for *N* samples, *W* denotes an *N* × *c* matrix of covariates, such as age and principal components (PCs) of ancestry, *γ* represents the corresponding fixed effects (including an intercept term), *X* is the *N* × *M* genotype matrix, *β* denotes the random effect sizes for each SNP, and *e* denotes random residuals so that *g* = *X β* and *e* represent the random additive genetic and environmental components, respectively. We simplify the notation by assuming that the term used to represent covariates *Wγ* has been regressed out from the phenotype *Y* and that *Y* is standardized to have zero mean and unit variance. We assume that 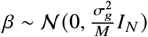 and 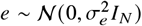, where 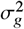 and 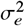 capture the genetic and environmental variances and *I*_*N*_ is the *N* × *N* identity matrix. We therefore have 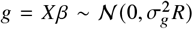, where *R* = *X X*^⊤^ denotes the *N* × *N* genetic relatedness matrix (GRM). Assuming that *g* and *e* are uncorrelated, the sample variance-covariance is given by 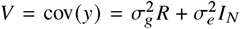 . In addition, *M* should be viewed as a general scaling factor to ensure Tr *R* = Tr *I*_*N*_ = *N*, so that 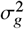 and 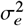 are on the same scale. This coincides with the number of SNPs *M* under the assumptions that genotypes have been standardized and are in Hardy–Weinberg equilibrium. Under this setting, each SNP contributes equally to the overall phenotypic variance, which is equivalent to choosing *α* = −1 in the model introduced in [73], where the *j* -th variant with frequency *f* _*j*_ has effect size *β*_j_ with variance 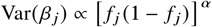 .

### 2.2. RHE algorithm

We describe the Randomized Haseman–Elston (RHE) regression algorithm [84], upon which ARG-RHE is based. The goal is to estimate narrow-sense heritability, the proportion of phenotypic variance attributable to additive genetic effects, defined as 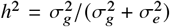 [78]. Assuming a single genetic component, the phenotype is modeled as *Y* = *g* + *e*, where 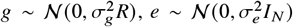, and the GRM is defined as *R* = *X X*^⊤^/*M*, with *X* ∈ℝ^*N* ×*M*^ the matrix of standardized genotypes. The moment-based estimators 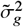 and 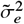 are obtained by solving the system:

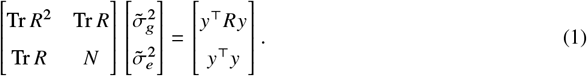

To avoid forming *R*, RHE uses Hutchinson’s stochastic trace estimator [24], where traces such as Tr *R* are approximated using *B* random vectors 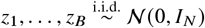,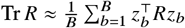. As detailed in [84],the resulting computational cost is 𝒪 (*NMB*), dominated by matrix-vector multiplications, and the memory cost is 𝒪 (*N* +*B*), assuming a streaming implementation that avoids storing the GRM or the genotype matrix. In contrast, standard restricted maximum likelihood (REML) methods such as GCTA [87, 85] involve operations on dense *N* × *N* GRMs, which scale at least quadratically with *N* and require 𝒪 (*N*^2^) memory. An extension of this approach, RHE-mc [62], generalizes the model to multiple variance components.

We further extended this approach to estimate heritability across *P* phenotypes in parallel with minimal additional computational cost. This is achieved in three steps: (1) compute *Y*^⊤^*RY* for each phenotype; (2) estimate Tr *R* and Tr *R*^2^ using shared random vectors; and (3) solve *P* linear systems using the same precomputed trace estimates. The overall complexity is dominated by terms proportional to *N M* (*B* + *P*), which can be substantially more efficient than analyzing each phenotype independently, where the cost scales as 𝒪(*NMBP*).As discussed in the Supplementary Information, additional care is required when phenotypes have differing patterns of missingness. Further implementation details, including extensions to the multi-component setting, are provided in the Supplementary Information.

### 2.3. ARG-matrix multiplication and ARG-RHE

A wide range of statistical and population genetics analyses, including RHE, can be framed in terms of multiplying a genotype matrix *X* by other matrices or vectors. When the mutations encoded in *X* are represented within an ARG, the structure of the graph can be leveraged to significantly reduce computational costs. This strategy, which we refer to as ARG-matrix multiplication (see also [27]), relies on graph traversal algorithms that efficiently propagate information across the nodes and edges of the ARG. We summarize the key components of this approach below and provide further details in the Supplementary Information.

The ARG is a graph representing the genealogical history of a set of homologous genomes, where nodes correspond to the haplotypes of samples or their ancestors, edges represent the transmission of genetic material, and mutations are annotated along these edges. It can be efficiently represented in memory using graph-based data structures and traversed using algorithms that compute properties of its internal nodes, edges, and mutations. Such representations and algorithms were originally developed as part of genome simulators, including ARGON [59, 61] and msprime [29, 4]. These simulators use graph traversal to sample mutations in a memory-efficient manner [59, 29], and similar strategies have been applied to compute ARG summaries such as identity-by-descent (IBD) haplotypes [59] and other statistics [66]. As detailed in the Supplementary Information, these algorithms leverage memoization, allowing properties of nodes or mutations, such as the set or number of descendants or carriers, to be computed efficiently by reusing results from nearby nodes or mutations.

Our approach for ARG-matrix multiplication builds on these ideas to efficiently compute matrix products of the form *U* · *X* (left multiplication) and *X* · *U* (right multiplication), where *X* is the *n* × *p* genotype matrix for *n* individuals and *p* mutations in the ARG, and *U* is an input matrix of compatible dimensions. Due to linkage disequilibrium (LD), the set of carriers of one mutation often overlaps significantly with carriers of other nearby mutations. In such cases, matrix-vector products can be computed by reusing results from related mutation vectors, enabling computations that scale sublinearly with respect to the number of samples, depending on the extent of LD present in the data.

We implemented the RHE algorithm from Section 2.2 using this approach, which we refer to as ARG-RHE. ARG-RHE leverages ARG-matrix multiplication routines to efficiently perform numerical computations involving repeated matrix–vector products, such as Hutchinson’s trace estimator and computing the leading eigenspace of the GRM *R* ∝ *X X*^⊤^ for significance testing (see Section 2.4). ARG-RHE, as well as other ARG-based analyses that can be built on this framework, can be applied either to the existing mutations present on an inferred ARG or to newly resampled mutations, as described in [90]. When newly resampled mutations are used, ARG-RHE captures the effects of variants likely present at the sequencing level but not directly observed in the data used to infer the ARG. This provides a computationally efficient alternative to the approach implemented in [90], where mutations were randomly resampled from an inferred ARG, written to disk, and explicitly used to construct ARG-GRMs for downstream analysis with GCTA. Such explicit construction limited those analyses to relatively small simulated samples.

In [90], the ARG was also used to perform large-scale LMM analyses of the UK Biobank dataset. Due to computational constraints, however, these analyses relied on the BOLT-LMM algorithm to implicitly construct GRMs from common SNP array variants, using the ARG only to test for association with unobserved variation. While ARG-RHE and our applications to large-scale data focuses on heritability estimation, ARG-matrix multiplication can also be leveraged to accelerate additional ARG-based LMM analyses. For instance, polygenic prediction and association testing require solving linear systems of the form *V x* = *Y* using conjugate gradients [45], where 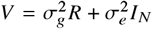 . In the Supplementary Information, we describe an efficient implementation of an ARG-based polygenic prediction and association testing approach that leverages ARG-matrix multiplication operations, as previously described in [27].

### 2.4. Efficient ARG-based variance component association testing

Here, we propose another extension so that, in addition to obtaining an estimate of the variance explained by mutations that may be generated using the ARG within a region, we can also efficiently assess whether this estimate is statistically significant, thereby performing variance component association testing.

For ARG-based association testing in this paper, we focus on the case of a single genetic variance component and a phenotype *y* standardized to unit variance, solving equation (1) while maintaining that Tr *R* = *N* leads to a closed-form expression for the heritability estimator

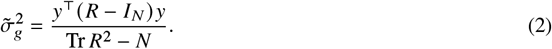

We consider the case where *R* is a local GRM, constructed by resampling mutations within the ARG in a region of interest. To test for association with the phenotype *Y*, we focus on the distribution of the estimand 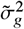 under the null hypothesis 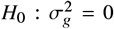, where 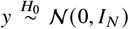 . The random part in the numerator of equation (2) gives the test statistic *T* := *Y*^⊤^*RY*. After eigen-decomposing *R* = Σ^⊤^ΛΣ, where Λ = diag(*λ*_1_, …, *λ*_*N*_) holds the eigenvalues of *R* in decreasing order and columns of Σ store the corresponding eigenvectors, we have

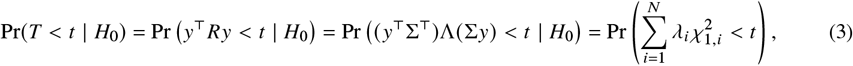

where *Y* ∼𝒩 (0, *I*_*N*_) is invariant under the action of Σ, and 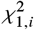 are i.i.d. chi-squared variables with one degree of freedom. This test statistic coincides with the score-based test adopted in methods such as the sequence kernel association test (SKAT) [83], which is instead derived by considering the derivative of the restricted likelihood at 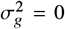, although SKAT has a different default weighting of variants based on their allele frequencies. We note that, while it is also possible to efficiently estimate the standard error of 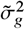 [84], using a Wald test on *H*_0_ requires *T* to be asymptotically normally distributed. However, since *T* in equation (3) has significant skew and kurtosis, convergence to a normal distribution depends on the number of non-zero *λ*_*i*_. This is bounded by the number of variants in a gene region and is usually not sufficient to apply large-sample asymptotics [76]. Hence, to obtain a tail probability that is precise enough for association testing but also computationally efficient, we follow the strategy recently proposed in fastSKAT [46], which relies on a mixture of moment-based and saddle-point approximations. More precisely, we first use ARG-matrix multiplication within a randomized SVD algorithm [68, 23] (also adopted to compute principal components and eigenvalues in PLINK2 [11]) to estimate *λ*_1_, …, *λ*_*L*_ as the top-*L* eigenvalues of Λ. We then approximate the right-hand side of equation (3) using a truncated sum of *L* + 1 chi-squared variables, which take the form

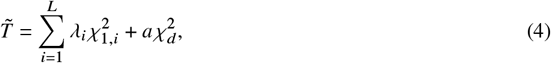

Where

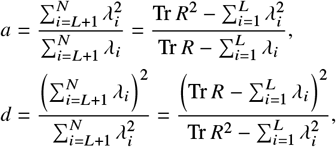

are scalars that satisfy the moment-matching conditions under the null 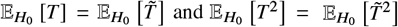 We estimate these quantities using the leading eigenvalues and Hutchinson’s estimator for the trace of powers of the GRM *R*. Next, we estimate association *p*-values via a less accurate but fast moment-based estimation approach [43]. If the *p*-value is below a specified threshold, which we set to 0.01, we proceed to estimate the tail probability more accurately through a saddle-point approximation technique [37]. This approach is designed to avoid the numerical instability of saddle-point approximation near the mean of the distribution. As done in the computation of point estimates for heritability, we use ARG-matrix multiplication to directly perform these computations within the ARG.

### 2.5. Simulation experiments

We evaluated ARG-RHE by testing its scalability, the accuracy of its *h*^2^ and *p*-value estimates, and its power when applied to perform variance component association analysis.

To work with Monte Carlo ARG-GRMs (i.e., ARG-GRMs estimated using mutations randomly resampled from the inferred ARG at high rate, see [90]), we discarded the original mutations used to infer the ARG and resampled a new set at a rate of 10^−6^ for 10-kb windows and 10^−8^ for 10-Mb windows. This yielded a genotype matrix *X*, which we zero centered and normalized by [ *f* (1 − *f*)] ^*α*/2^, where *f* denotes the allele frequency.With this approach, the ARG-GRM can be written as *R* = *X X*^⊤^/*M*, where *M* is a normalizing factor such that Tr *R* = *N*, and used to calculate *p*-values of the test statistic *T* = *Y*^⊤^*RY*. In experiments involving GCTA, we explicitly formed the ARG-GRM after writing these mutations to disk, as done in [90]. When running the ARG-RHE algorithm, the mutations remained stored in the ARG and were used to obtain heritability estimates and assess significance. We used 200 random vectors to compute Hutchinson’s trace estimator and the top 200 leading eigenvalues obtained from the randomized SVD. All tests were performed using Intel(R) Xeon(R) Gold 6240R 2.40GHz CPUs.

#### 2.5.1. Heritability estimation accuracy

To assess the accuracy of the heritability estimates, we generated ARGs containing 20,000 individuals under a constant effective population size *N*_*e*_ = 50,000 and recombination rate *ρ* = 10^−8^, spanning 10 kb and 10 Mb gene windows. We then generated heritable traits with *h*^2^ ranging from 0 to 0.8. We varied polygenicity from 10% to 80% and the MAF-dependency parameter *α* from −1 to 0.5 (Supplementary Table 4). We generated 100 independent simulations for each combination of parameters. When estimating *h*^2^ using ARG-RHE, we assumed the correct *α* used to generate the phenotypes.

We also performed experiments in which we estimated 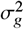 using time-stratified GRMs. For these analyses, we simulated 10-Mb windows on 20,000 individuals as above. We additionally partitioned the resampled genotypes into 6 GRMs containing an equal number of mutations based on their age quantiles, and jointly estimated the 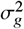 across time bins using multi-component ARG-RHE. We generated heritable traits with *h*^2^ =0.4 and effect sizes *α* ranging from −1 to −0.25, and formed the ARG-GRMs using a fixed *α* = −1.

#### 2.5.2. Computational scalability

We simulated 10-kb and 5-Mb regions with up to 1,000,000 diploid individuals under a demographic model inferred from the GBR population in the 1,000 Genomes Project dataset [75]. These simulations were used to assess the scalability of the ARG-matrix multiplication routine and its applications, using resampled mutations generated at specified rates. We compared the total CPU time required with that of other baselines, including NumPy and PLINK2, applied to the same set of resampled mutations. To assess computational efficiency, we again simulated 10-kb gene regions with up to 500,000 individuals and compared the costs of running ARG-RHE to the use of GCTA to estimate heritability with an explicitly constructed ARG-GRM, as done in [90] and [40]. When comparing the use of GCTA to the ARG-RHE algorithm, we used the same set of resampled mutations to explicitly construct the ARG-GRM using GCTA (v1.94.1) or to run the ARG-RHE algorithm, which is implemented in the `arg-needle-lib` library (see URLs). We measured the computational time required to run GCTA on pre-computed ARG-GRMs, excluding the time required to construct such ARG-GRMs, and compared with the time required to compute heritability and test for significance using ARG-RHE. Unlike ARG-RHE, GCTA does not support analyzing multiple phenotypes in parallel, so in these benchmarks, we tested ARG-RHE with a single phenotype, as well as with 50 random phenotypes processed in parallel.

#### 2.5.3. p-value calibration and statistical power

We performed additional simulations to verify that variance component testing using ARG-RHE results in sufficiently calibrated *p*-values. To this end, we obtained exact quantiles for the test statistic *T* at *p*-values ranging from 0.05 to 10^−8^ for simulations consisting of 5,000 individuals. We used Davies’ method [12], obtaining a complete SVD of *R* to verify that the randomized approach used by ARG-RHE produces accurate *p*-values for a range of significance levels. For each quantile, we tested 20 independently generated ARGs each with 5 sets of random trace estimator seeds, totalling 100 estimators at each *p*-value threshold. We also verified the empirical null distribution of the *p*-values by testing 50 randomly generated non-heritable traits against the same inferred gene-based ARGs from 337,464 UK Biobank individuals used in our real data analysis.

We evaluated the statistical power of ARG-RHE under a range of setups to assess the complementarity of its gene-based association signals to alternative approaches such as ACAT-V [44] and MAGMA [13] applied to imputed or exome-sequenced data. We simulated ARGs covering a 20-Mb region under a demographic model inferred from the GBR population in the 1,000 Genomes Project dataset [75], including up to 200,000 diploid individuals, along with an additional 10% used as an imputation reference panel. Sequencing variants were simulated by sampling mutations from the true ARG at a rate *μ* = 10^−8^. For samples not in the reference panel, we filtered mutations to match the allele frequency spectrum of UK Biobank SNP array data, following the approach in [90]. These SNP variants were also used as input to Beagle v5.5 for imputation and to Threads v0.1 [20] to construct the inferred ARG, using default parameters.

We generated synthetic phenotypes by selecting the central 10-kb segment of the ARG as the gene region and randomly choosing causal variants from the simulated sequencing variants within this region, sampling their effects as *β* ∼ℕ (0, [ *f* (1− *f*)] ^*α*^) with *α* = −1. We added additional noise to achieve a trait heritability of *h*^2^ = 0.01. To avoid simulating rare variants with unrealistically large effect sizes, we restricted causal variants to those with minor allele count (MAC) ≥ 5. To simulate causal variants that are not included in the imputation panel, we generated mutations on the same ARG using a different random seed. To model different causal allele frequency patterns, we sampled causal variants either from recent time (i.e., up to 1,000 generations) or uniformly across the entire ARG. After evaluating the effect of varying the resampling rate on ARG-RHE’s power (see Supplementary Information and Supplementary Figures S2(a) and S2(b)), we chose 10^−6^ as the default mutation sampling rate on inferred ARGs. For each parameter combination, we generated 20 sets of ARGs and sequencing variants, each with 20 independently drawn sets of causal mutations and phenotypes. Statistical power was assessed at *p* = 2.5 × 10^−6^, corresponding to a Bonferroni-corrected threshold assuming 20,000 gene regions and a family-wise error rate of 0.05.

We also compared the statistical power of ARG-RHE testing, which uses a moment-based approach, to that of GCTA, which relies on a more accurate but less scalable likelihood-based approach. We ran GCTA using two sets of variants for the GRM. In one case, we computed the local GRM using the actual set of sequencing variants used to simulate the phenotype. This simulation provides an upper bound to the statistical power that can be achieved using ARG-RHE, which relies on a different set of resampled mutations. In the other case, we explicitly constructed a Monte Carlo ARG-GRM, using the same set of resampled mutations used by ARG-RHE as input GRM for GCTA to test for association. This simulation is meant to assess the extent to which the approximate randomized approach of ARG-RHE results in decreased association power compared to GCTA’s likelihood-based approach. We again assessed the statistical power at *p* = 2.5 × 10^−6^, and performed 100 repetitions for each combination of parameters.

#### 2.6. Analysis of UK Biobank data

We applied ARG-RHE to perform variance component association testing using an ARG inferred for 337,464 unrelated white British individuals from the UK Biobank dataset [10], using a single variance component for resampled variants within a gene-based window. The ARG was inferred using Threads [21] (v0.1) applied with default parameters to 798,103 genotyped variants, each with less than 10% missing values, which were computationally phased using Beagle (v5.1) [9]. We tested for association between 21,159 protein-coding and non-coding RNA regions from the HUGO Gene Nomenclature Committee (HGNC) table on the University of California, Santa Cruz (UCSC) genome browser (see URLs) and 52 quantitative blood cell traits, including 27 blood cell indices and 25 blood biochemistry marker levels. The lists of tested regions and traits are provided in Supplementary Tables 1 and 2.

We compared the results obtained using ARG-RHE to those obtained by applying the same variance component association test of equation (4) to genotype dosages imputed from the Haplotype Reference Consortium (HRC) reference panel, as performed in [10]. We included HRC-imputed variants with MAC ≥ 5,INFO score ≥ 0.3, missingness ≤ 10%, and Hardy-Weinberg equilibrium *p* ≥ 10^−15^, leading to a total of 54,356,141 variants tested for standard single-variant association testing using Regenie [49]. We refer to these single-variant analyses of HRC-imputed data as HRC-SV. We also computed the test statistic of equation (4) using 22,460,173 HRC-imputed variants that overlap a gene region, which we refer to as HRC-RHE. Finally, we also performed ACAT-V gene-based testing [44] by combining HRC-SV *p*-values from a gene region using uniform weights. We set the genome-wide significance threshold to be 0.05/21159 ≈ 2.36 × 10^−6^ for gene-based tests, and 10^−9^ for single-variant tests.

We compared the signals from both gene-based and single-variant tests by grouping *p*-values from the tested regions into approximately independent LD blocks, using LDetect [6]. We then checked whether each block-trait pair contains genome-wide significant signals. We also considered a simple joint test that combines ARG-RHE and HRC-RHE: for each gene region, we took the smaller *p*-value *p*_min_ from the two methods and treated them as independent tests by comparing 2*p*_min_ to the same genome-wide significant threshold. We used a similar method for combining ARG-RHE, HRC-RHE, and ACAT-V results. When running ARG-RHE, we used *L* = 200 leading eigenvalues in estimating the tail distribution and *B* = 200 random vectors in Hutchinson’s trace estimator.

To preprocess the analyzed traits, we followed the approach of [3]. We first stratified samples based on sex and menopause status and applied a rank-inverse-normal transformation (RINT) to the adjusted phenotype values. We regressed out covariates, which included age, age squared, alcohol use, smoking status, height, body mass index, assessment centre, genotyping array, and 20 principal components. We then applied RINT a second time and merged all strata. Missing phenotypes were mean-imputed as 0. To verify that the top 20 PCs can account for most of the population structure, we performed simulations in which heritable traits were simulated using HRC-imputed variants on odd chromosomes and residualized using the leading 20 PCs. We then tested for associations with regions on the even chromosomes and observed no significant deviation from the null (see Supplementary Information and Supplementary Figure S3). We also verified that mean imputation at 0 does not inflate the type I error rate (see Supplementary Information and Supplementary Figure S4).

We verified the detected associations by checking whether they were previously reported on the Open Targets platform (version 22.10) [18, 53]. Specifically, for each significant gene-trait association, we used Open Targets’ locus-to-gene pipeline to check for associated variants from previous studies involving the same gene. We considered an LD block-trait pair as previously detected if the LD block contained at least one such associated gene for the trait in Open Targets. For this analysis, we matched our 52 quantitative traits to 495 traits with closely related descriptions available on Open Targets (Supplementary Table 3).

We additionally validated our genome-wide significant associations against signals detected using Genebass [28], a phenome-wide association study based on exome sequencing data from a larger UK Biobank cohort. For each gene-trait pair, we obtained summary statistics under four variant masks: synonymous, high-confidence predicted loss-of-function (pLoF), low-confidence pLoF and in-frame indels (missense LC), and the combined missense LC & pLoF variants. Each mask was analyzed with SAIGE-GENE [92] under three association methods: burden, SKAT, and SKAT-O. We grouped these gene-based exome summary statistics into our approximately independent LD blocks. To quantify enrichment, we computed the fold-increase in probability that an LD block contains a genome-wide significant exome signal, conditional on it being significantly associated by one of the tested methods (ARG-RHE, HRC-RHE, or ACAT-V). We compared this against the baseline probability of association, calculated across all LD blocks and irrespective of their significance under any method. We estimated the standard errors by resampling LD block–trait pairs with replacement. Finally, we focused on the subset of 27 blood cell count traits, and evaluated the detected gene associations using the GENE2FUNC module from FUMA [79] to annotate their expression levels in whole blood (see Supplementary Information).

## 3. Results

### 3.1. Scalability of ARG-matrix multiplication

We first evaluated the computational efficiency of ARG-matrix multiplication, which underlies ARG-RHE, by comparing its runtime to that of two optimized numerical libraries, NumPy and SciPy, for simulated ARGs and genotype data covering a 5-Mb genomic region (Methods). In these experiments, traditional matrix-based routines exhibited approximately linear scaling with sample size, whereas ARG-matrix multiplication scaled sublinearly, achieving speed-ups of over 15× at sample sizes approaching 500,000 individuals (Figure 1(a,b)). We observed that the performance benefit of ARG-matrix multiplication varies with the size of the analyzed region, with larger regions yielding greater speed-ups due to increased long-range LD among variants. In smaller regions (e.g., 10-kb windows; Figure 1(c), Supplementary Figure S1), ARG-matrix multiplication still provided a 5–10× speed-up relative to traditional routines, although the runtime scaled approximately linearly with sample size. Similar gains were observed in more complex numerical routines, such as truncated eigendecomposition of the GRM used to compute test statistics within ARG-RHE. Using ARG-matrix multiplication to compute the top 20 principal components (PCs), we observed a ∼15× speed-up for 1,000,000 samples relative to an optimized implementation of the same randomized SVD algorithm in the PLINK2 software package [11] (Figure 1(d)). Finally, we confirmed that the generality of ARG-matrix multiplication enables additional speed-ups in downstream analyses beyond those implemented in ARG-RHE. For example, in Figure 1(e), we used ARG-matrix multiplication to compute a key quantity related to the best linear unbiased predictor (BLUP) of a trait by evaluating products involving the inverse GRM using the conjugate gradients algorithm of [45], and observed a ∼9× speed-up for 200,000 individuals compared to a NumPy-based implementation. Additional details, including the use of this approach to perform scalable ARG-based LMM association, as previously described in [27], are provided in the Supplementary Information.

**Figure 1:**
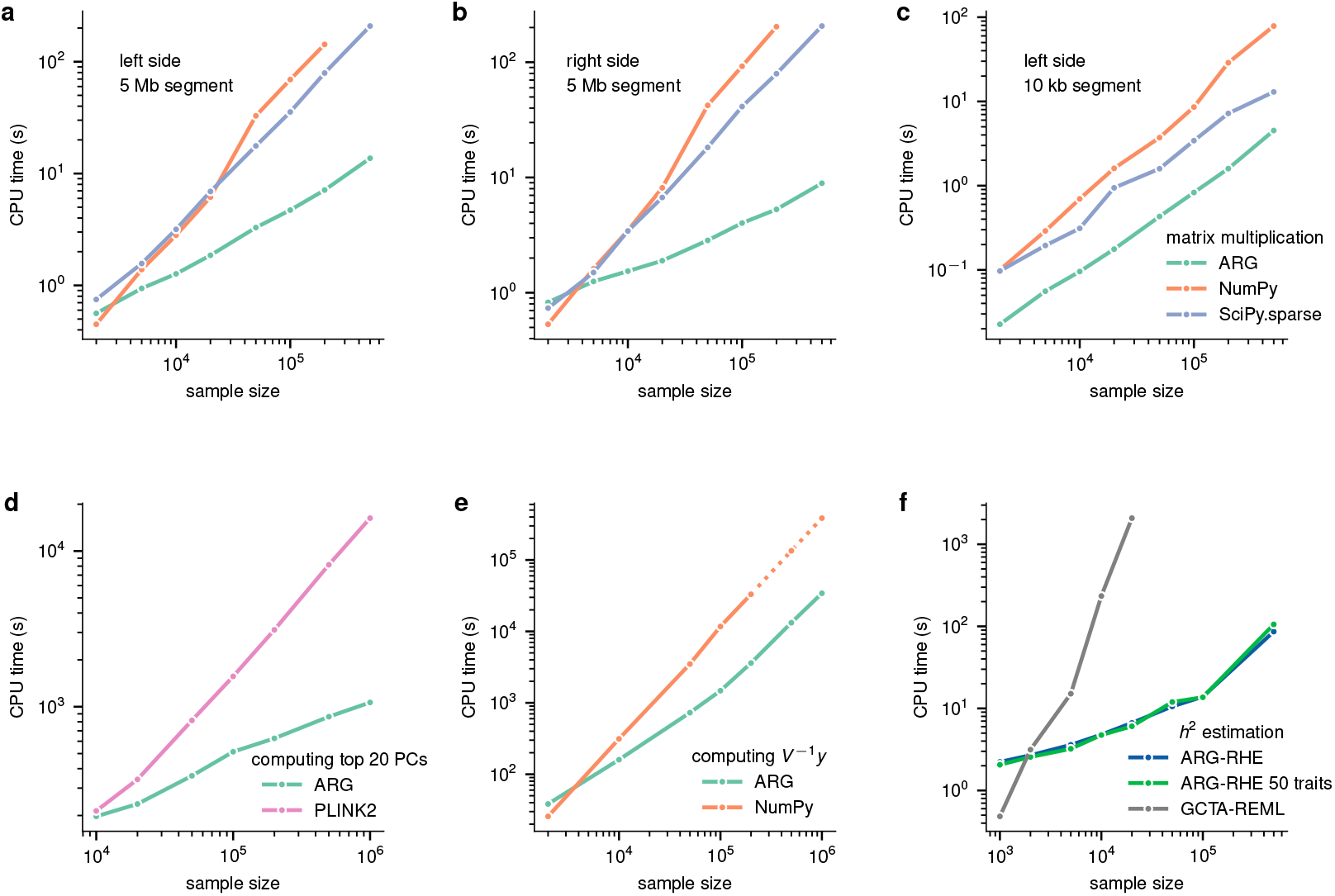
Scalability of ARG matrix multiplication algorithm. (a, b, c) CPU time taken to calculate genotype matrix-matrix product using various methods, based on the average runtime from 10 replicates. In (a, c), left side multiplication computes *UX*, and in (b) right side multiplication computes *XU* where *U* is a random input matrix with one dimension implied by the *n* × *p* genotype *X*, and the other being 100. In (a, b), ARGs are simulated under the GBR demographic model and cover a 5Mb region, with mutations resampled on the ARG at rate 10^−8^. In (c), ARGs are simulated under the GBR demographic model and cover a 10kb region, with mutations resampled on the ARG at rate 10^−6^. (d) CPU time taken to calculate top 20 principal components using various methods. ARGs are simulated under the GBR demographic model and cover a 30Mb region. Mutations are resampled on the ARG at rate 10^−8^, and filtered to have MAF at least 1%. 4 power iterations are used in the ARG-based randomized PCA scheme. (e) CPU time required to compute *V* ^−1^ *Y*, used to obtain the BLUP or to residualize the phenotype *Y* for LMM association testing, where 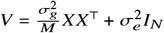 . This is computed using conjugate gradient iterations implemented with either ARG-matrix multiplication (see Supplementary Information, Note 3.3; see also [27]) or NumPy. (f) CPU time and required for testing a single trait or 50 traits in parallel on a simulated 10 kb window for sample sizes up to 500,000 individuals. Both ARG-RHE and GCTA-REML used random mutations generated from the ARG at a rate of *μ* = 10^−6^.

#### 3.2. Scalability, accuracy, and power of ARG-RHE

We next performed extensive simulations to verify the scalability, accuracy, *p*-value calibration, and statistical power of variance component analyses using ARG-RHE. We first verified the scalability of ARG-RHE under large sample sizes by performing association in up to 500,000 individuals. We compared ARG-RHE to the approach of [90] and [40], which is based on first computing an ARG-GRM and then providing it to GCTA to estimate heritability using restricted maximum likelihood (GCTA-REML, Methods). The results of these analyses are shown in Figure 1(f). As expected, GCTA-REML, which involves manipulating dense ARG-GRM matrices containing *N*^2^ entries for *N* individuals, required substantially more computation and memory than ARG-RHE. As an example, testing an ARG region for association using 20,000 individuals with GCTA-REML required ∼300× more time and ∼24× more memory. The gap widened as the sample size increased, with an extrapolated difference of four orders of magnitude for an analysis involving 500,000 individuals. In addition, we verified that our extension of the RHE algorithm allows for efficiently estimating heritability across multiple traits in parallel, requiring negligible additional computation to analyze 50 traits compared to a single trait, as shown in Figure 1(f).

We evaluated the accuracy of using ARG-RHE to infer heritability across 96 genetic architectures simulated using different parameters (see Methods). The results of these analyses are shown in Figure 2 and Supplementary Tables 4 and 5. Applied to 10 Mb gene windows, ARG-RHE yielded accurate estimates, with an estimated relative bias ranging from −3.6% to 1.0%, which is consistent with results observed when applying the RHE algorithm to genotype data [62]. On smaller 10 kb gene windows with simulated *h*^2^ up to 0.02, ARG-RHE yielded slightly conservative estimates of heritability, with a relative bias ranging from −2.6% to −19.4%, depending on the *α* parameter, which governs the relationship between allele frequency and effect sizes [73, 70]. A small downward bias in ARG-based heritability estimates may also be observed if the mutation rate used when resampling mutations to work with Monte Carlo ARG-GRMs is not sufficiently large [90].

**Figure 2:**
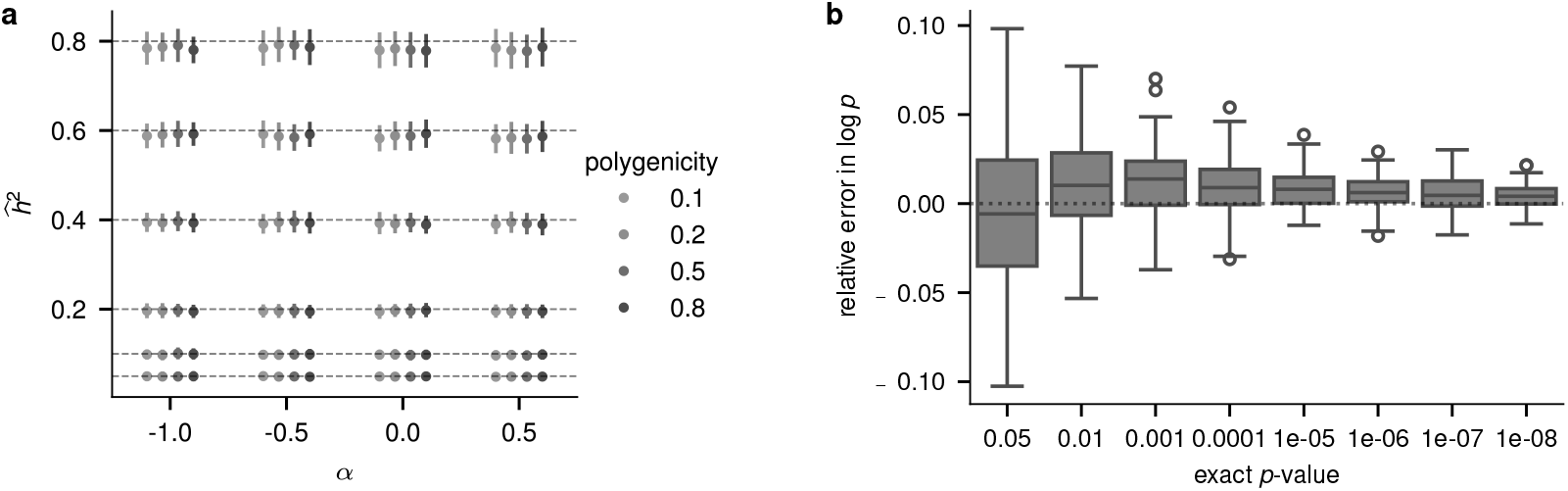
Accuracy of estimated heritability 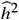 and *p*-values using ARG-RHE. (a) Estimated heritability 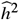 over a range of polygenicity levels, true *h*^2^ (shown as dashed lines), and MAF-dependency parameter *α*. Heritability was estimated for a 10-Mb region and 20,000 samples, resampling mutations at a rate of *μ* = 10^−8^. Error bars show standard deviations estimated using 100 independent repetitions. (b) Distribution of the relative error in estimated *p*-values ( − log_10_ *p*_approx_ + log_10_ *p*_exact_)/( − log_10_ *p*_exact_) corresponding to exact *p*-value quantiles, using *L* = 200 leading eigenvalues across 100 random samples. *p*-values were estimated for a 10-kb region and 5,000 samples, resampling mutations at a rate of *μ* = 10^−6^.

In addition to representing information on the genotypes of the analyzed samples, the ARG contains information on the age of the analyzed variants and can therefore be used to assess the phenotypic contribution of variants originating at different time scales. These analyses could allow estimating the strength of natural selection on variants linked to the trait [48, 65, 36, 33] or help quantify the relative phenotypic contribution of variation that is yet to be sequenced. To verify that ARG-RHE allows studying time-stratified variance components, we simulated phenotypic data using MAF-dependent architectures under different values of *α*, corresponding to different effect size distributions for variants of different allele frequencies (see Methods). Due to the link between the frequency and age of an allele [34, 19], these simulated architectures also imply different effect size distributions for variants of different ages, with larger negative values of *α* leading to larger effect sizes for rarer and more recent variants. These trends were captured by ARG-RHE when we applied it to jointly estimate multiple 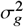 values using implicit time-stratified GRMs (see Methods), as shown in Supplementary Figure S5.

We next assessed the calibration of *p*-values obtained using ARG-RHE when performing variance component association testing. We compared the *p*-values from ARG-RHE, which are based on a fast randomized algorithm (see Methods), to the exact *p*-values obtained using Davies’ method. To this end, we assessed the relative error between exact and approximate log-*p*-values, (− log_10_ *p*_approx_ +log_10_ *p*_exact_)/(− log_10_ *p*_exact_), shown in Figure 2(b). Consistent with previous applications of this class of randomized approaches [46], we found a small inflation in the mean estimated log *p*-value, particularly at more extreme *p*-values, which corresponds to the exact critical values being estimated to be positioned around 0.93 times their nominal tail quantile. To verify that this inflation is unlikely to meaningfully impact our UKB analysis, we used the same set of inferred gene-based ARGs, simulated 50 non-heritable traits, and tested for association, observing 5 gene-trait pairs with *p*-values that exceeded the Bonferroni-corrected threshold. This is a modest increase over the expected number of false positives under the null (0.05 × 50 = 2.5, S.D. ∼1.9), and represents a negligible fraction of the total associations detected in real data, described below.

Next, we evaluated the ability of ARG-RHE, applied to SNP array data, to detect association signals complementary to those identified by other set-based methods, such as ACAT-V [44] and MAGMA [13], when applied to imputed or sequenced variants. Results of these analyses are shown in Figure 3. To reflect varying genetic architectures and imputation accuracy, we simulated different scenarios by varying the age of causal variants and the extent to which these variants were polymorphic in the imputation reference panels (Methods). In a scenario where 10% of causal variants were included in the imputation panel, comparable to recent estimates for the proportion of exome sequencing variants accurately imputed using the HRC panel [2, 72], combining ARG-RHE with other tests yielded substantial power gains over imputation alone. The complementarity observed in these simulations was consistent with our analyses based on HRC-imputed UK Biobank data. As expected, in cases where all causal variants were accurately imputed, as in analyses based on directly sequenced data, ARG-RHE did not provide additional power. However, across a range of settings where imputation was incomplete, the combination of ARG-RHE and imputation-based methods consistently yielded the highest power, suggesting that ARG-RHE can effectively complement genotype imputation, particularly in scenarios where causal variants are poorly tagged by reference panels.

**Figure 3:**
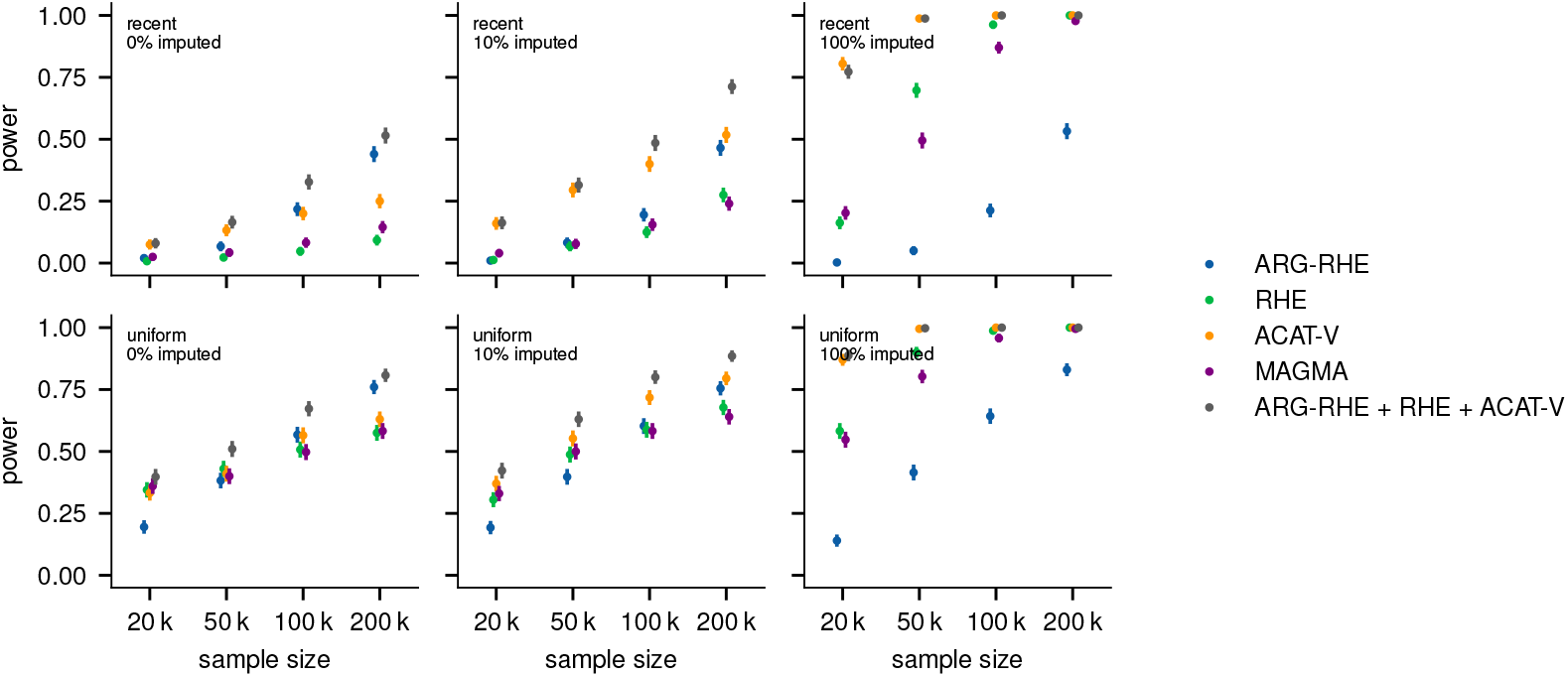
Statistical power of ARG-RHE with a varying proportion of causal variants included as imputation targets. We computed association power using a 10 kb gene window under polygenicity level *g* = 0.05 (fraction of causal variants among all sequencing variants simulated at rate 10^−8^), local heritability *h*^2^ = 0.01 (fraction of phenotypic variance explained by causal variants), and significance threshold *s* = 2.5 × 10^−6^. Causal variants are either sampled within the last 1,000 generations (recent) or uniformly throughout the entire genealogy (uniform), with 0%, 10% and 100% of them being included as imputation targets. Error bars show standard errors estimated from 20 independently simulated ARGs each with 20 simulated traits.

Finally, we assessed the extent to which using fast randomized algorithms results in a drop in association power compared to GCTA’s likelihood-based approach. To this end, we compared ARG-RHE and GCTA-REML applied to the same set of mutations resampled from the ARG, which are distinct from the underlying causal variants (see Methods). Results for varying polygenicity and significance settings (see Methods) are shown in Supplementary Figure S6. At a significance threshold of 2.5×10^−6^, ARG-RHE achieved a statistical power of around 60% relative to GCTA-REML for *h*^2^ up to 0.02. Previous studies have observed a similar difference in statistical power between kernel-based score tests, such as in SKAT, compared to the likelihood ratio test used in GCTA [41]. However, due to its computational demands, GCTA cannot be applied to largescale datasets for this analysis

Overall, ARG-RHE provided substantial gains in computational speed and memory requirements compared to GCTA-REML, while providing good accuracy and well-calibrated *p*-values. The high computational scalability of this approach comes at the cost of a reduction in association power compared to exact likelihood-based strategies, which is consistent with previous applications of randomized approaches to variance component association testing. However, this decrease in power is offset by the possibility of applying ARG-RHE to datasets containing orders of magnitude more individuals.

#### 3.3. Analysis of a UK Biobank ARG

We applied ARG-RHE to an ARG comprising 337,464 unrelated white British individuals from the UK Biobank dataset, which was inferred using 798,103 genotyped variants. We leveraged this ARG to test for association between 21,159 protein-coding and non-coding RNA regions and 52 quantitative blood cell counts and serum biomarker level traits (see Methods). We compared the results of ARG-RHE to those obtained by applying the same variance component association test to ∼22 million variants imputed using the HRC reference panel (HRC-RHE). We also compared to associations detected by running Regenie to perform single-variant testing using the same set of HRC-imputed variants (HRC-SV), and to using the ACAT-V algorithm to combine single-variant association statistics from imputed variants within gene regions (see Methods). Results of these analyses are shown in Figure 4. The computing resources required to apply HRC-RHE and ARG-RHE to the UK Biobank were lower than those required by highly optimized GWAS approaches. Applying HRC-RHE to test 52 traits across 21,159 regions spanning ∼22 million imputed variants required 299.6 CPU hours, ARG-RHE applied to the UK Biobank ARG required 601.2 CPU hours, and applying Regenie to individually test the imputed variants required 730.8 CPU hours, without including step 1 of the analysis (see Figure 4(a)). We first considered the sets of approximately independent genomic regions [6] in which gene-trait associations were detected using each method, reported in Figure 4(b). Taken individually, running ARG-RHE, which relies on an ARG inferred using only genotype array markers, identified 10,374 unique locus-trait associations, while HRC-RHE and ACAT-V, which rely on imputed data, identified 12,162 and 13,720, respectively. Consistent with recent analyses [90, 40], however, we found association signals detected using the ARG to complement signals detected using imputation alone. Among the signals identified using ARG-RHE, 1,858 (18%) were not found by HRC-RHE, 1,376 (13%) were not found by ACAT-V, 1,770 (17%) were not found by HRC-SV, and 1,054 (10%) were not found by either HRC-RHE or ACAT-V (Figure 4(c) and Supplementary Figure S7). Performing a joint test using both imputed variants and the inferred ARG (see Methods) yielded an increase in the number of associations. Combining ARG-RHE with HRC-RHE led to 13,190 associations, 8.5% more than only using HRC-RHE; ARG-RHE with ACAT-V led to 14,401 associations, 5% more than ACAT-V alone; while combining all three tests led to 14,831 associations, 21.9% and 8.1% more than only using HRC-RHE and ACAT-V, respectively. The full list of all genome-wide significant associations found by ARG-RHE and estimated heritability 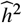 are provided in Supplementary Table 6.

**Figure 4:**
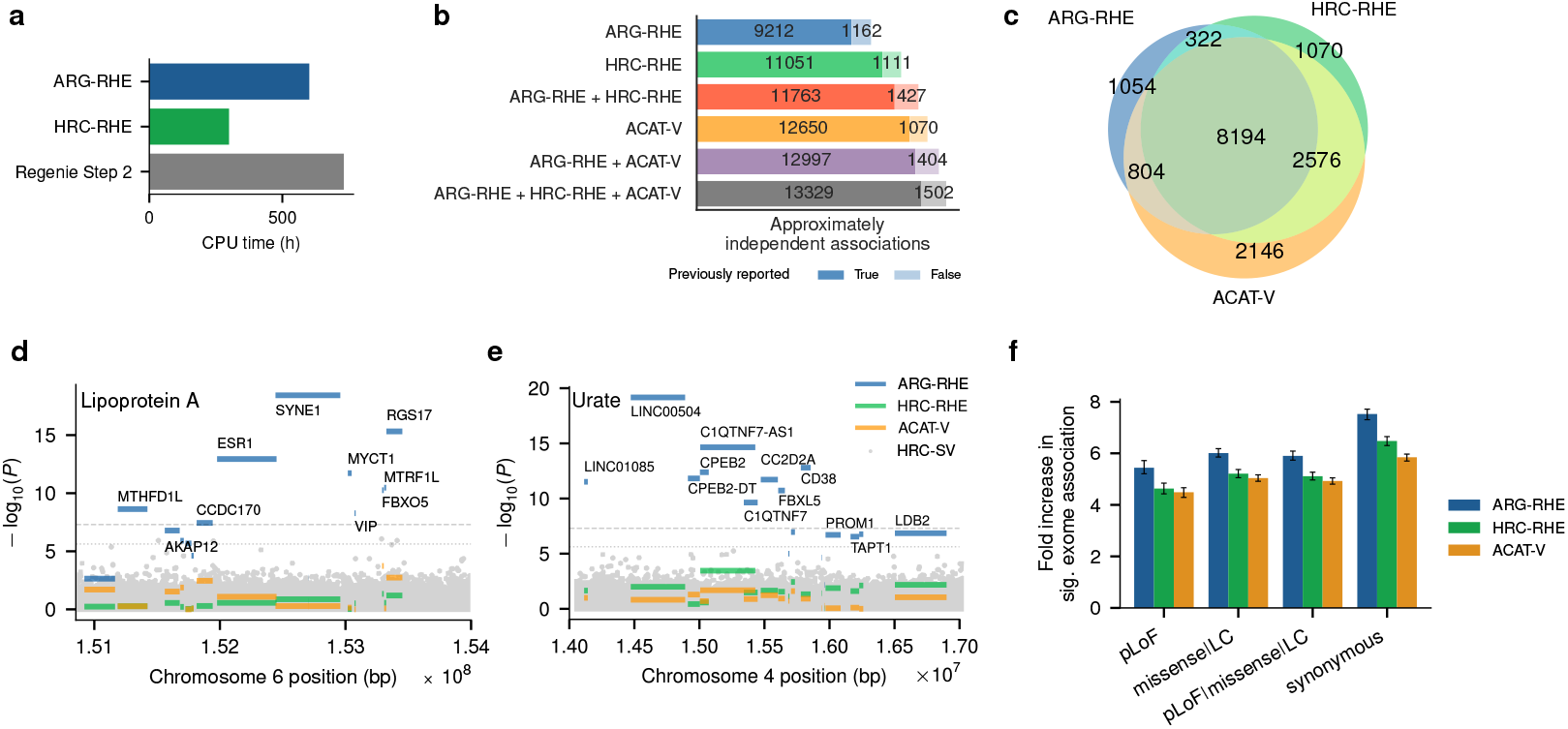
UK Biobank association analyses. (a) Total CPU time taken to test 52 traits for 337,464 unrelated white British individuals from the UK Biobank dataset. Both the ARG-RHE and HRC-RHE methods were run on a single thread, while Regenie was run using 12 cores; we report the total CPU time across all cores. (b) Total number of LD blocks containing genome-wide significant gene-trait associations detected using different methods, including their combination using joint tests. We annotate the counts by whether the significant LD block-trait pair contains known gene-trait associations reported on the Open Targets platform (see URLs). (c) Overlap in LD blocks containing gene-trait associations detected using ARG-RHE applied to ARGs inferred from SNP array data, or HRC-RHE, ACAT-V applied to HRC-imputed variants. (d) Manhattan plot for a genomic region containing genes, including SYNE1 and ESR1, associated using ARG-RHE to lipoprotein A levels; and similarly (e) for a genomic region containing LINC00504 non-coding RNA, CPEB2, and CD38 genes, associated with serum urate levels. Dotted lines represent the genome-wide significance threshold for gene-based tests. Dashed lines represent the genome-wide significance threshold for HRC-SV single-variant tests. (f) Concentration of significant exome gene-based associations with 52 blood traits after conditioning on the LD block having a genome-wide significant association in 337,464 unrelated white British individuals. Four annotation sets are used to filter the exome variants to be included in the gene-based SKAT-O test (see Methods; pLoF: high confidence predicted loss-of-function variants, missense LC: low-confidence pLoF variants and in-frame indels, missense LC & pLoF which is the combination of the previous two masks, and synonymous). Error bars show bootstrapped standard error in the estimated mean.

We sought to validate the associations detected using ARG-RHE by looking for overlap with associations from other studies reported in the Open Targets database [58] (see URLs), which includes summary statistics from several single-variant and gene-based association studies, including analyses of large whole-exome or whole-genome sequencing datasets. Overall, we found 9,212 of the 10,374 regions associated with a trait by ARG-RHE to be reported on the Open Targets platform (see Methods). These include established signals found using ARG-RHE but not detected in our analysis of this subset of UK Biobank samples using HRC-RHE, HRC-SV, or ACAT-V, such as the association between RNF168 with red blood cell count [32], and the association of SLC4A4 with neutrophil count [3]. Of the remaining 1,162 signals that are not present in Open Targets, several have been linked to related traits in other studies. Examples include associations found on Chromosome 6 with lipoprotein A (Lp(a)), shown in Figure 4(d). Among the highly significant genes detected using ARG-RHE, SYNE1 has a reported association with Lp(a) levels in the Hutterites, a founder population of South Dakota [57]. Associations with ESR1 have been reported in [35], and the RGS17 gene was significantly associated using exome sequencing data [3]. Other signals uniquely identified using ARG-RHE in our analyses and not reported in Open Targets include loci previously linked to traits that are related to those we have found to be associated. An example is the associations found on Chromosome 4 with serum urate levels, shown in Figure 4(e). The LINC00504 non-coding RNA and the CPEB2 gene were found to be associated with gout in a recent study involving ∼2.6 million individuals [47] and the CD38 gene was found to be associated with gout in another study [80].

We also examined whether regions uniquely identified by ARG-RHE differ systematically from those uniquely identified by HRC-RHE or ACAT-V (Supplementary Table 7). On average, genes with associations detected by ARG-RHE were shorter, exhibited higher recombination rates, and had lower imputation INFO scores. While some gene-based signals may reflect LD with neighboring regions, these trends support the hypothesis that ARG-RHE is particularly effective in regions with reduced imputation quality, where alternative methods may have reduced power. We further compared the associations detected by ARG-RHE and other methods to gene-based summary statistics derived from rare variant tests in UK Biobank exome sequencing data [28]. Figures 4(f) and S8 show the enrichment of exome-based signals across LD blocks found significantly associated with 52 blood phenotypes using each method. LD blocks containing signals detected by ARG-RHE using SNP-based ARGs were more likely to overlap with rare exome-based associations than those identified by HRC-RHE or ACAT-V, suggesting that ARG-RHE may be more sensitive to rare variant signals that are poorly imputed. Finally, using FUMA’s GENE2FUNC pipeline [79], we verified that genes associated with blood cell count traits by ARG-RHE showed enrichment for expression in whole blood in GTEx [77], comparable to those identified by HRC-RHE and ACAT-V (see Supplementary Information and Supplementary Figure S9).

Overall, the ARG-RHE algorithm was scalable enough to be applied to a genome-wide ARG inferred for 337,464 UK Biobank individuals using only genotyping array variants, and allowed detecting associations that complemented those that would be identified using the same variance component test applied to HRC-imputed data alone. Signals uniquely detected using ARG-RHE in these analyses were often implicated by studies that involved different cohorts or closely related traits. Further applications of ARG-RHE to non-European UK Biobank subgroups are described in [21].

## 4. Discussion

We developed ARG-RHE, an approach that uses ARG-based computation and builds on recent work on moment-based estimation of narrow-sense heritability [62] and on randomized algorithms to approximate tail probabilities [83, 46] to enable highly scalable heritability estimation and variance component association testing using an ARG. In simulations, ARG-RHE was several orders of magnitude faster than the use of explicitly computed ARG-GRMs to estimate heritability [90] and to perform variance component association testing [40]. We verified the accuracy, calibration, and association power of ARG-RHE in simulations, and confirmed its ability to identify complementary association signals, particularly when causal variants are poorly tagged by genotyped or imputed markers. Finally, we applied ARG-RHE to an ARG inferred for 337,464 UK Biobank individuals using only genotyping array variants, detecting associations that complement signals identified using the same gene-based variance component test on ∼22 million HRC-imputed variants or single-variant linear mixed model testing applied to ∼54 million HRC-imputed variants.

Our study contributes to a growing body of work leveraging inferred genome-wide genealogies for the analysis of complex traits [90, 69, 40]. One key advantage of using the ARG in this setting is that it provides a route to account for genomic variants that cannot be observed or accurately imputed. This is particularly the case in populations that are poorly represented in sequencing reference panels used for genotype imputation [90, 40], as also evidenced by further applying ARG-RHE to non-European UK Biobank individuals in [21]. Compared to testing individual edges of an inferred ARG, as done in [90], the use of a variance component test improves association power by pooling the effects of multiple variants and reducing the statistical burden by reducing the number of performed tests [40]. Our work substantially improves the computational efficiency of ARG-based variance component analyses of heritability and association, making this approach easy to apply to significantly larger samples.

Although our UK Biobank applications focused on association testing, our simulations show that ARG-RHE is scalable enough to infer genome-wide, rather than local, narrow-sense heritability. By leveraging information on allele ages contained in the ARG, it may also be used to gain insights into time-stratified phenotypic variance components. Natural selection acting on variants linked to polygenic traits often leads to larger per-allele effect sizes for variants of lower frequencies and younger ages [48, 65, 36, 33], with stronger selection leading to a larger overall phenotypic contribution from rare and recent variation [89, 70]. ARG-based, time-stratified analysis may therefore quantify the strength of selection on variants across different time strata, complementing models that link selection and allele frequencies [64, 73, 89, 70], as well as previous time-stratified approaches limited to higher frequency variation [17, 16]. These analyses may provide insights into the fraction of phenotypic variance explained by rare and recent alleles, which may not yet have been sequenced and are more likely to be population specific. However, we caution that obtaining unbiased heritability estimates is more challenging than testing for significance, as it imposes additional requirements on the accuracy of the inferred ARG. Current ARG benchmarks suggest that the accuracy of inferred topologies and branch lengths depends on factors such as marker density, recombination-to-mutation rates, and time scale [90, 20], making robust time-stratified heritability analysis an interesting direction for future work.

We highlight several additional avenues for future development and limitations of this work. First, ARG-RHE currently only supports quantitative traits, but it could be extended to support the association of rare variants with binary traits that have low case/control ratios [91, 49, 25]. Second, ARG-RHE implements a simple form of over-dispersion testing. Future work may involve implementing different strategies for weighting and pruning markers to improve power under various assumptions about the trait’s architecture [44, 7, 94,13, 92, 5]. Third, the multi-trait extension of the RHE-mc algorithm, which we use to reduce computational cost, relies on a single estimate of *H* and *b* in equation (1), under the assumption that the intersection of individuals with non-missing phenotype data is large and that GRM estimation is not biased by missingness. This approach may be extended to obtain different *H* and *b* estimates for each trait. Fourth, as with other gene-based or single-variant association approaches, signals detected using ARG-RHE may reflect LD with variants in neighboring genes, and additional downstream analyses are required to more precisely localize causal variation. Finally, beyond heritability analysis, ARG-matrix multiplication can accelerate computations for other tasks, such as computing polygenic predictors (BLUP; Figure 1(e)) or performing mixed-model association (see Supplementary Information), as also described in [27]. Extending these methods to additional analyses and applying them to population-scale datasets is a promising direction for future work. Despite current limitations and areas of future work, the computational strategies developed here provide a highly scalable framework for leveraging inferred ARGs to complement genotype imputation in mixed-model analyses.

## 5. URLs

`arg-needle-lib` library, implementing ARG-RHE, https://palamaralab.github.io/software/argneedle

GENIE library, implementing multi-trait RHE-mc for genotype data, https://github.com/sriramlab/GENIE

Open Targets database, https://genetics.opentargets.org

UCSC genome browser, https://genome.ucsc.edu/cgi-bin/hgTables

## 7. Acknowledgments

This research was conducted using the UK Biobank Resource (application 43206). We thank the participants of the UK Biobank project. This work was supported by the EPSRC Centre for Doctoral Training in Health Data Science (EP/S02428X/1, to J.Z.); ERC Starting Grant 850869 (to P.F.P. and G.K.); EPSRC and MRC grant EP/L016044/1 (to G.K.); the Keble College de Breyne Clarendon Scholarship (to A.F.G.); Wellcome Trust grant 222336/Z/21/Z (to A.F.G.); the Clarendon Scholarship (to H.L.); Wellcome Trust Studentship (to H.L.); NSF CAREER 1943497 and R35GM153406 (to S.S.). Computation used the Oxford Biomedical Research Computing (BMRC) facility, a joint development between the Wellcome Centre for Human Genetics and the Big Data Institute supported by Health Data Research UK and the NIHR Oxford Biomedical Research Centre. Financial support was provided by the Wellcome Trust Core Award Grant Number 203141/Z/16/Z. The views expressed are those of the author(s) and not necessarily those of the NHS, the NIHR or the Department of Health.

## 8. Supplementary Figures

**Figure S1:**
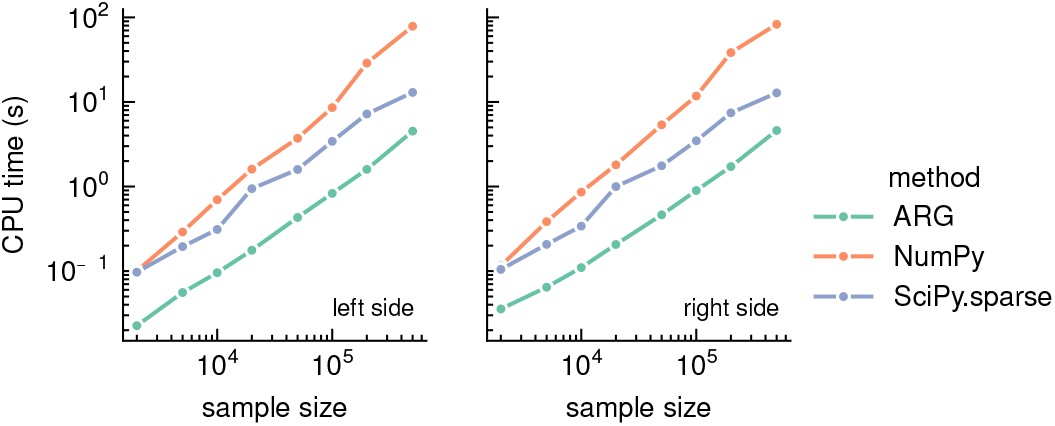
CPU time required to compute genotype matrix–matrix products using various methods, averaged over 10 replicates for a 10-kb region. “*n* side” refers to computing *UX*, and “ *p* side” refers to *XU*, where *U* is a random matrix with one dimension matching the genotype matrix *X* and the other set to 100. ARGs were simulated under the GBR demographic model, with mutations resampled at a rate of 10^−6^.

**Figure S2:**
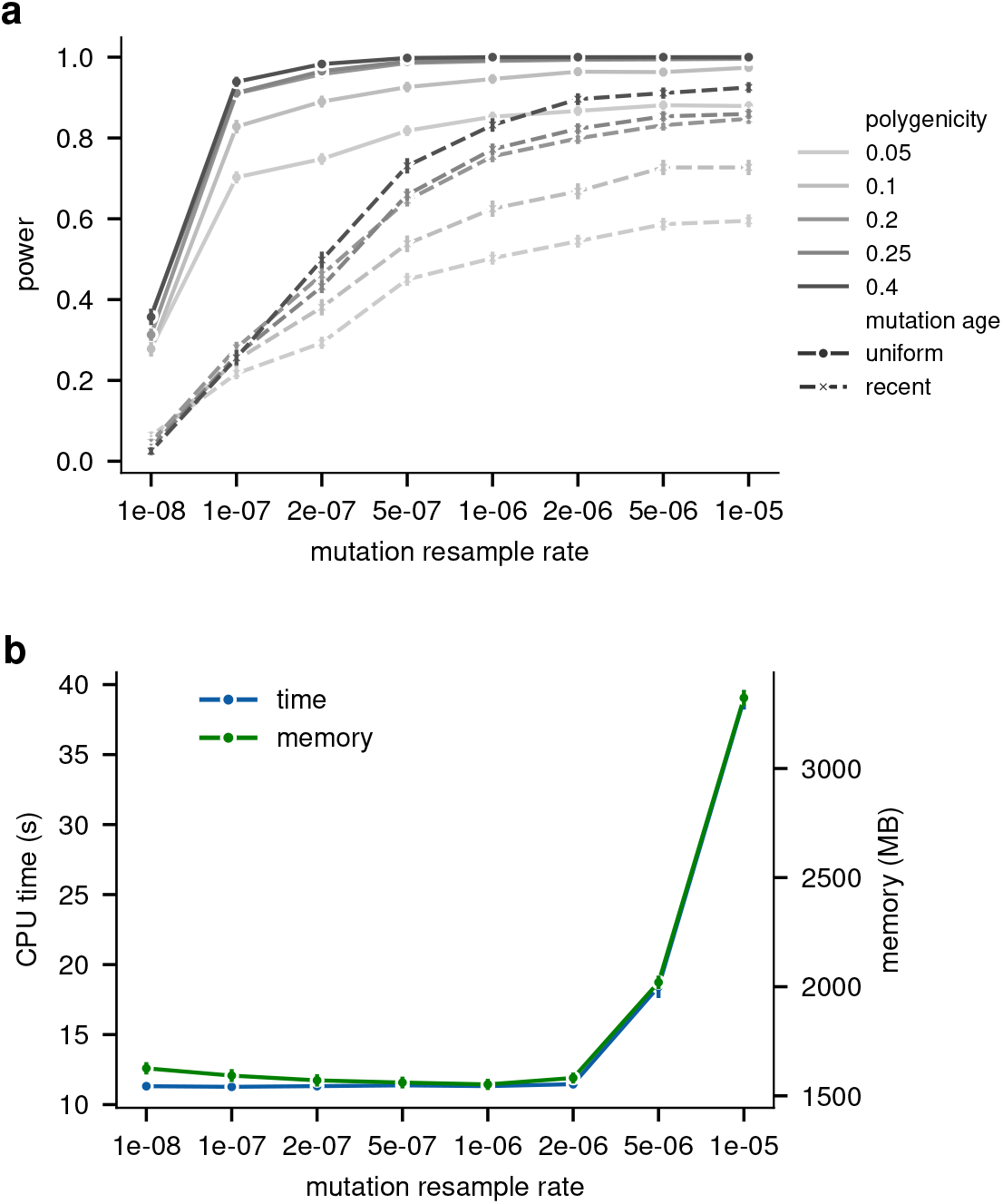
(a) Association power of ARG-RHE under different genetic architectures and increasing mutation resampling rates. Vertical bars indicate the standard error of the estimated mean power. (b) CPU time and memory usage of ARG-RHE as a function of mutation resampling rate. Vertical bars indicate the standard error of the estimated mean.

**Figure S3:**
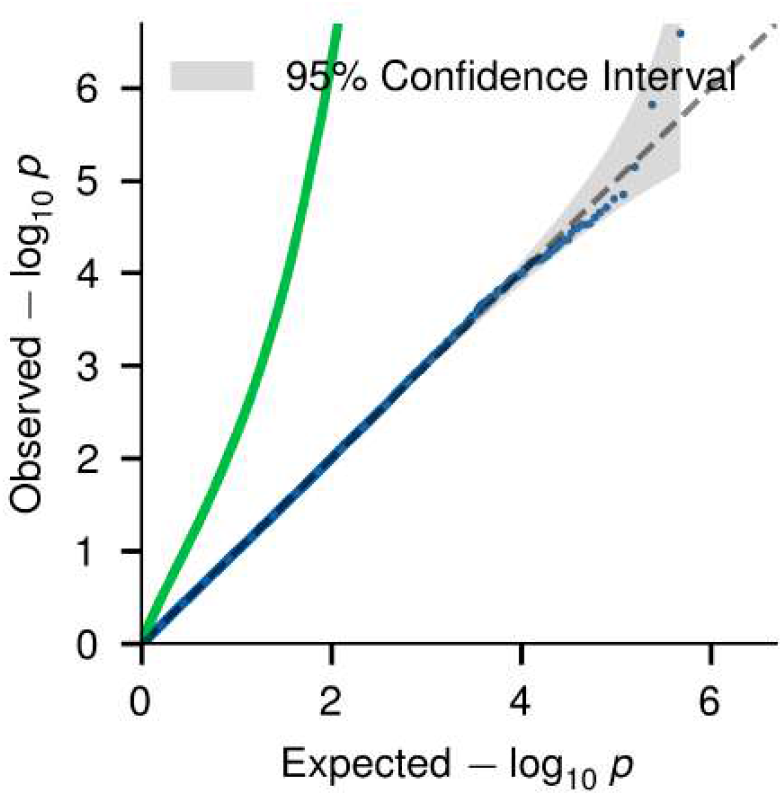
Quantile-quantile plot for − log_10_ *p* values from ARG-RHE using inferred ARGs testing for associations between genes on the even-numbered chromosomes (blue) and odd-numbered chromosomes (green) and 50 phenotypes, generated by causal variants located on the odd-numbered chromosomes only with *h*^2^ = 0.5, after controlling for the top 20 PCs.

**Figure S4:**
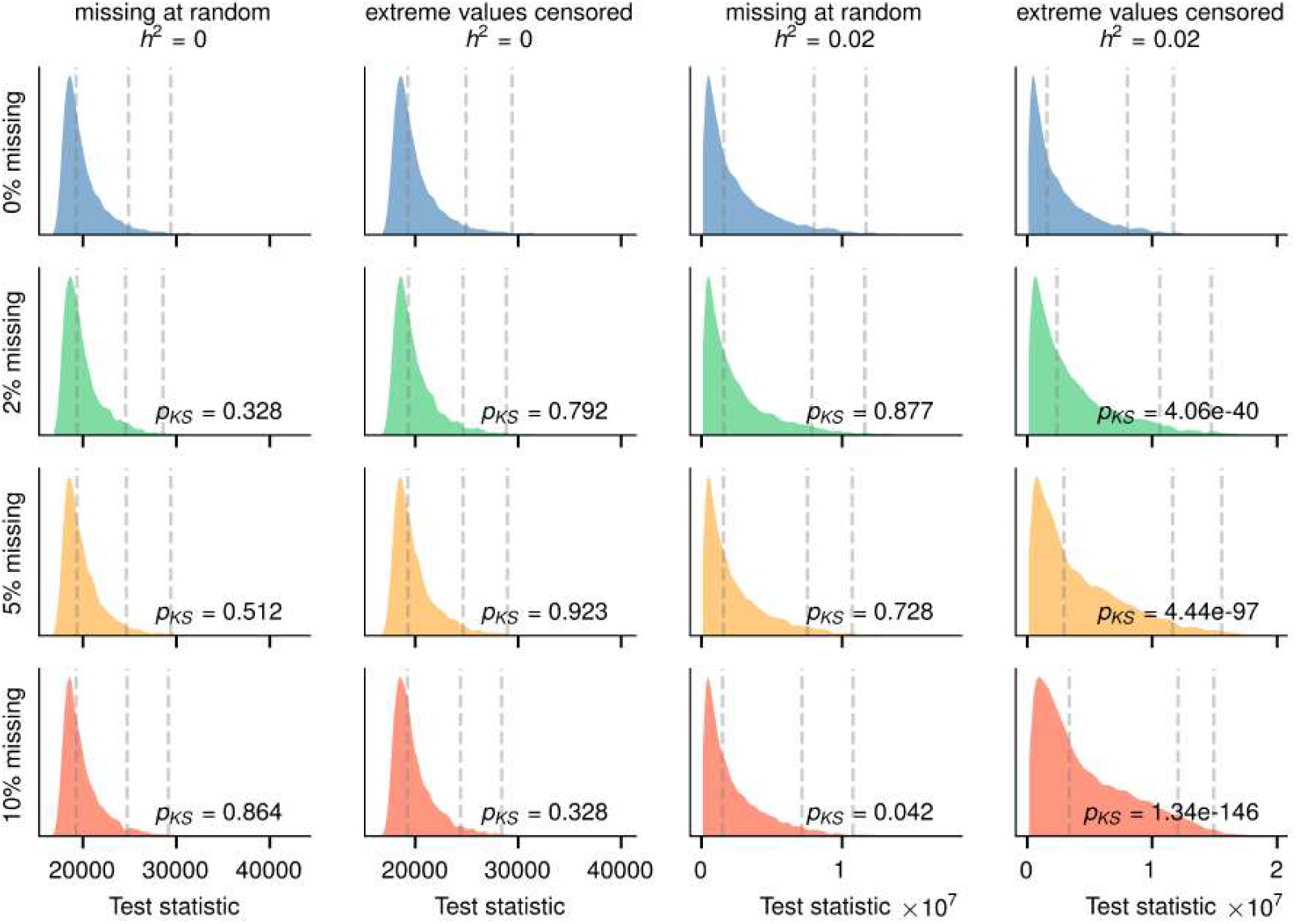
The distribution of test statistic under the null and alternative hypotheses with different missingness mechanisms. The dashed lines are the empirical 50th, 95th and 99th percentiles. *p*_*KS*_ is the *p*-value from Kolmogorov-Smirnov test comparing the test statistic distribution to the corresponding non-missing case.

**Figure S5:**
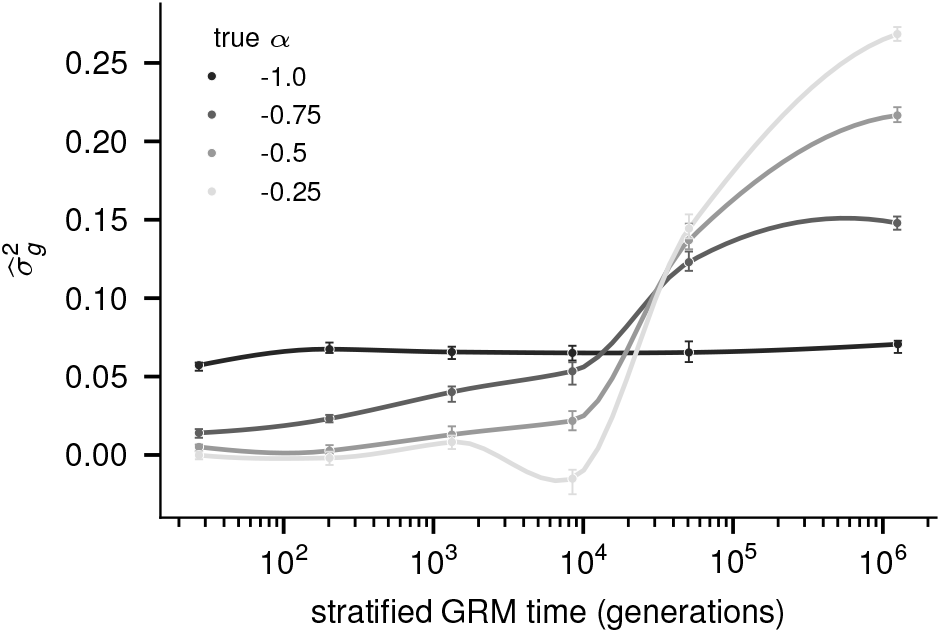
Joint estimates of heritability using time-stratified GRMs. Analysis of a 10-Mb gene window for 20,000 individuals with a heritable phenotype (*h*^2^ = 0.4, polygenicity *g* = 0.5). Mutations resampled from the ARG are grouped into six time bins, each containing an equal number of mutations, and separate 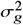 values are estimated for each bin. The true underlying MAF-dependency parameter *α*, which mimics the effect of negative selection by modulating the heritability contribution of variants by age, ranges from −1 to −0.25. All GRMs are constructed assuming *α* = −1. We report the median of the estimated 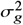 values across 500 replicates; error bars show 95% confidence intervals.

**Figure S6:**
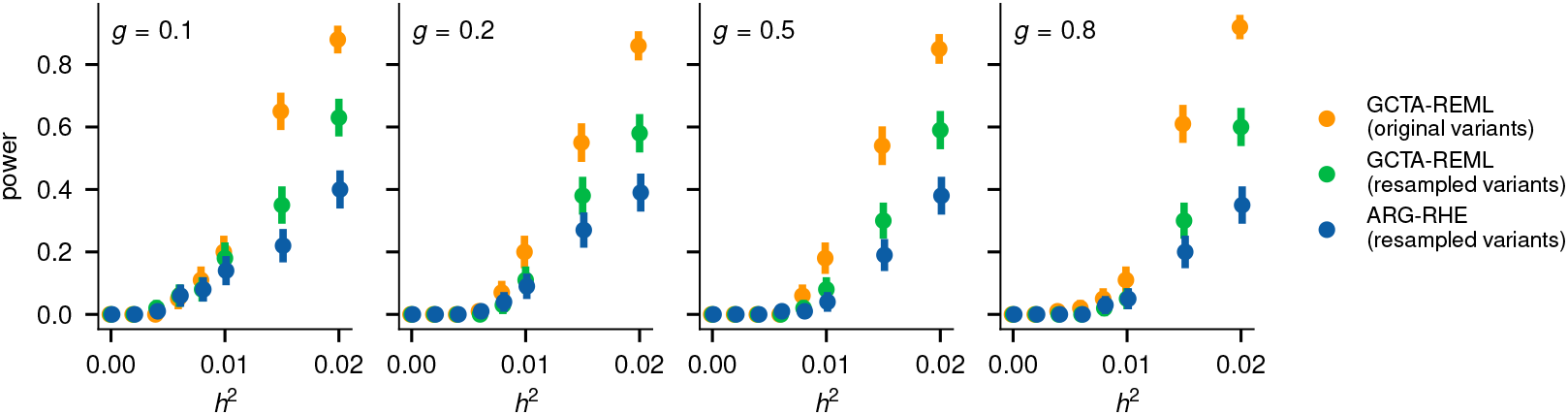
Statistical power of ARG-RHE. We computed power for a 10-kb gene window under different polygenicity levels *g* (fraction of causal variants among all sequencing variants simulated at rate 10^−8^), local heritability *h*^2^ (fraction of phenotypic variance explained by causal variants), and significance threshold 2.5 × 10^−6^. We performed variance component association using either GCTA-REML applied to all sequencing variants, GCTA-REML applied to variants resampled from the ARG at a rate of 10^−6^, and ARG-RHE applied to the same resampled variants. Error bars show standard errors estimated from 100 independent repetitions.

**Figure S7:**
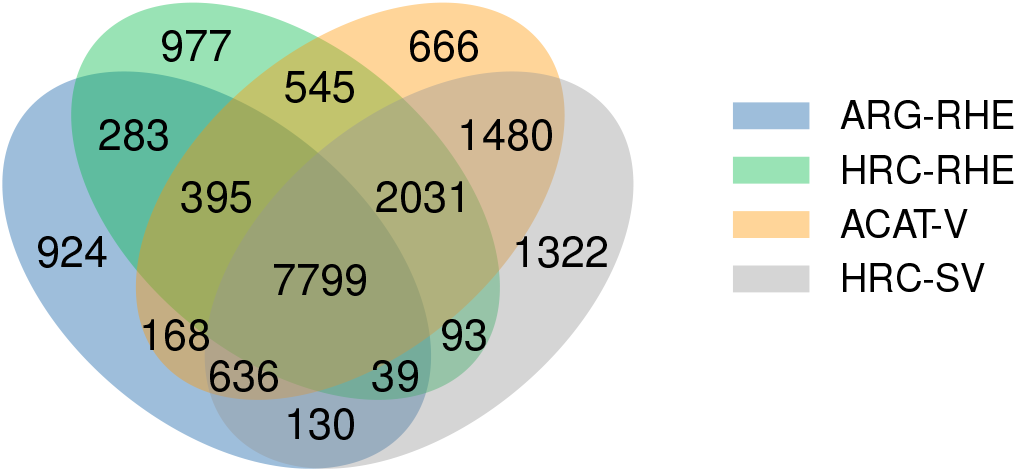
Overlap in LD blocks containing gene-trait associations detected using ARG-RHE and other methods. As in Figure 4(c)b, now including HRC-SV. ARG-RHE was applied to ARGs inferred using Threads from SNP array data, while HRC-RHE, ACAT-V, and HRC-SV were applied to HRC-imputed variants.

**Figure S8:**
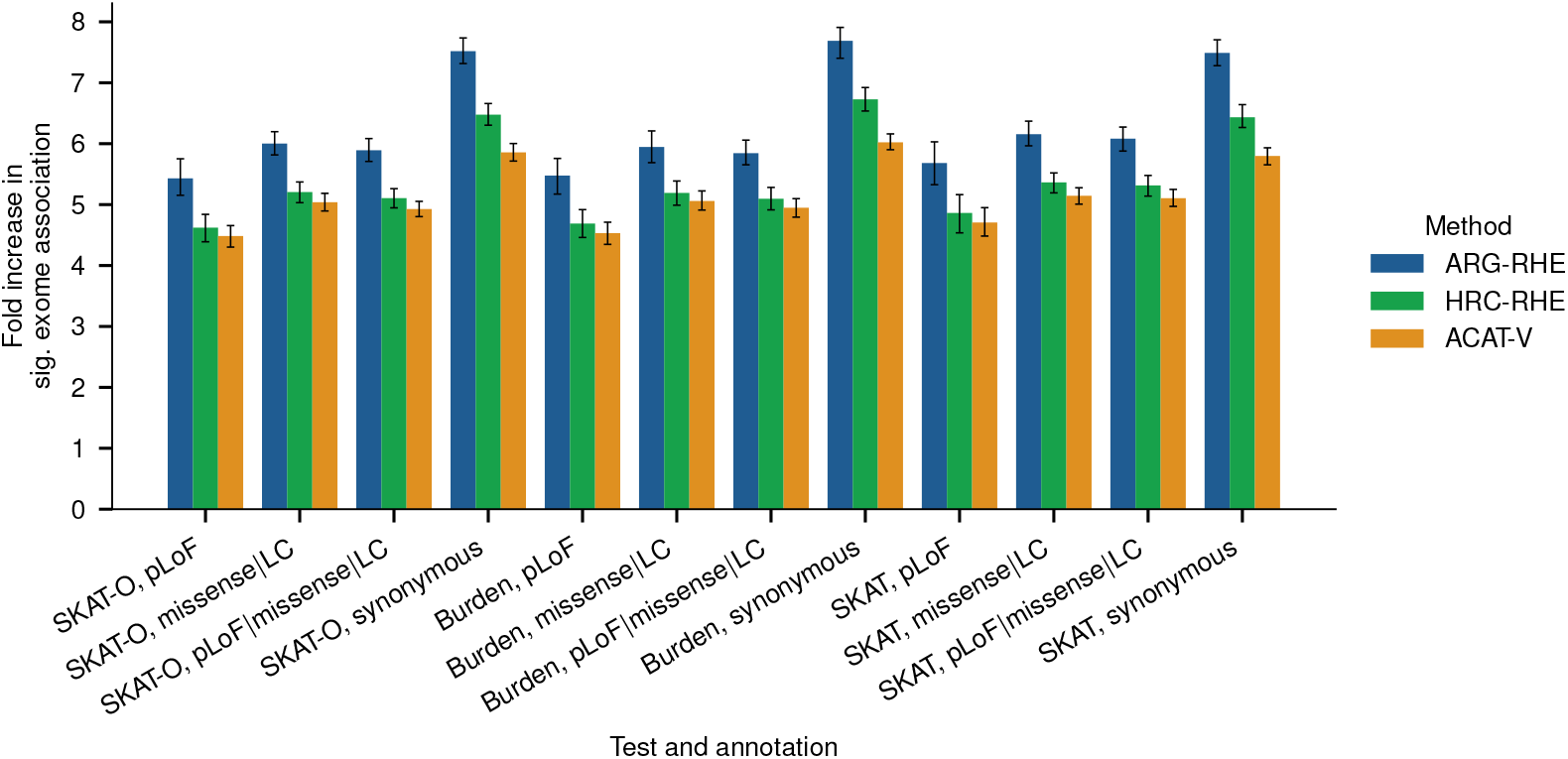
Concentration of significant gene-based exome associations for 52 blood traits, conditioned on the LD block having a genome-wide significant association in 337,464 unrelated white British individuals. Four annotation sets are used to filter exome variants for SKAT-O, burden, and SKAT tests (see Methods): pLoF (high-confidence predicted loss-of-function variants), missense LC (low-confidence pLoF and in-frame indels), missense LC & pLoF (combined), and synonymous variants. Error bars indicate bootstrapped standard errors of the estimated means.

**Figure S9:**
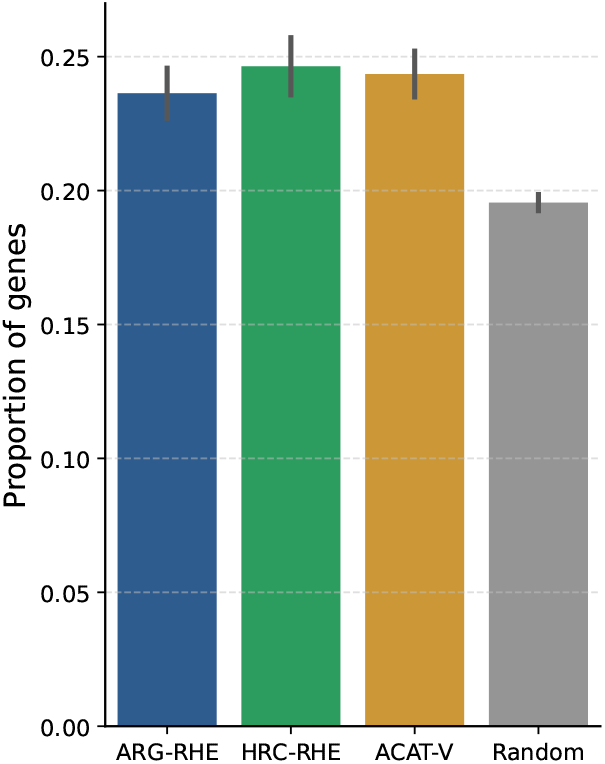
Gene expression relevance for associated genes. We report the fraction of genes significantly associated with blood-related phenotypes that are expressed in whole blood. Results are shown for ARG-RHE, HRC-RHE, and randomly sampled gene-phenotype pairs.

## Supplementary Information

## 1 Additional details on ARG-RHE

This section provides additional details on the multi-component RHE (RHE-mc) algorithm, its extension to handle multiple traits in parallel, the effects of phenotype missingness, and of resampling mutation rates.

### 1.1 Efficient heritability estimation across multiple traits

In the main text, we described the RHE approach using a single genetic variance component, which is also used by ARG-RHE for association testing. Here, we summarize how heritability can be partitioned across multiple components in the RHE-mc framework of [1], and outline how we extended RHE to handle multiple traits in parallel.

#### Multiple genetic components (RHE-mc)

RHE-mc [1] computes a moment-based estimator of heritability, allowing it to be partitioned across multiple variance components. This is achieved by extending the linear model for the phenotype *Y* = *g* + *e* to consider a decomposition of the genetic component *g* into *K* ≥ 1 components, *g*_1_, *g*_2_, …, *g*_*K*_, with *M*_1_, *M*_2_, …, *M*_*K*_ variants, respectively. Thus, the model becomes:

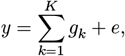

Where 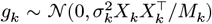, *X*_*k*_ is the *N* × *M*_*k*_ matrix of standardized genotypes for component *k*, 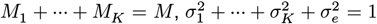, and 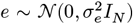 . Here, 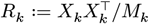 denotes the GRM for the *k*-th component. The moment-based estimators 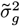 and 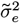 satisfy the normal equations:

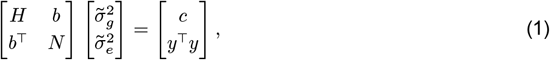

where 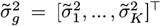, *H* is a *K* × *K* matrix with entries *H*_*k,ℓ*_ = Tr(*R*_*k*_ *R*_*ℓ*_), *b* is a *K*-vector with elements Tr(*R*_*k*_), and *c* is a *K*-vector with *c*_*k*_ = *Y*^T^*R*_*k*_*Y*. To avoid explicitly forming GRMs, RHE-mc estimates these traces using Hutchinson’s trace estimator [1–3] with random vectors. For genotype-based input, RHE-mc also leverages the Mailman algorithm to accelerate matrix operations. This is not used in ARG-RHE, which instead performs operations directly on the ARG.

#### Extension to multiple traits

We extended RHE to allow simultaneous estimation of heritability across *P* phenotypes with minimal additional computational cost. This extension, used in ARG-RHE, estimates trait-specific heritability independently across traits, but does not model genetic correlation between them. The procedure involves three main steps:

- For each of the *P* phenotypes, compute the vector *c* with elements *c*_*k*_ = *Y*^T^*R*_*k*_*Y* on the right-hand side of equation (1).
- Estimate the *K*^2^ elements of *H* using *B* shared random vectors.
- Invert *H* and solve *P* linear systems to obtain trait-specific variance component estimates.

The total computational complexity is 𝒪(*NM*(*B* + *P*) + *K*^2^*NB* + *K*^3^ + *P K*^2^), where the dominant term is typically 𝒪(*NM*(*B* + *P*)) and significantly smaller than the 𝒪(*NMBP*) cost incurred when analyzing each phenotype independently; the other terms reflect the cost of solving the normal equations for K components.

### 1.2 Effects of phenotype missingness

When estimating heritability or performing association testing across multiple traits in parallel, phenotype missingness poses a practical challenge. In the absence of missing data, the left-hand side (LHS) of the normal equations (equation (1)) is independent of the phenotype, enabling shared computation across traits. However, when phenotypes have differing patterns of missingness, the LHS becomes phenotype-specific, requiring separate computation of *H* and *b* for each trait using only the subset of individuals with observed values.

To maintain computational efficiency, our default implementation assumes that the intersection of non-missing samples across traits is sufficiently large. In this setting, we estimate *H* using the full genotype matrix, and impute missing phenotype values as zero after applying rank-inverse normal transformation (RINT). The resulting vectors are then re-normalized to have unit variance across all samples.

To justify this approach, consider the test statistic *T* = *Y*^T^*RY* for a phenotype vector *Y* and GRM *R*.Let 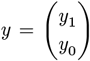 denote the phenotype values for *n*_1_ observed and *n*_0_ missing individuals, respectively, with *R* partitioned accordingly into blocks *R*_11_, *R*_10_, *R*_01_, *R*_00_. When missing values are imputed as zero after normalization, the test statistic becomes:

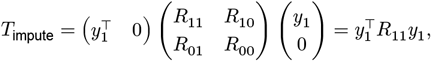

where *Y*_1_ is the mean-centered and rescaled vector ensuring unit variance across the observed samples.

Alternatively, one can exclude missing entries and compute:

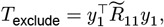

where 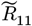 is the GRM recomputed using only the non-missing individuals, with allele frequencies estimated from that subset. Assuming missingness does not bias allele frequency estimation, we expect 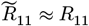, implying *T*_exclude_≈ *T*_impute ._

Under the null, the distribution of *T*_exclude_ is a weighted sum of *𝒳*^2^ random variables with weights given by the eigenvalues λ_*i*_ ∈ σ(*R*_11_), where σ(·) denotes the spectrum of a matrix:

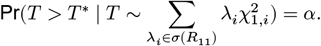

In contrast, *T*_impute_ is evaluated against a distribution that includes the full spectrum:

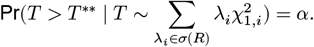

Since σ(*R*_11_) ⊂ σ(*R*), *T* ^∗∗^ > *T* ^∗^, making the imputation-based approach more conservative for type I error control.

To empirically assess the effects of missingness and this imputation scheme on ARG-RHE, we conducted simulations under two missingness mechanisms: (1) missing at random, and (2) missing values preferentially affecting phenotypes with the largest absolute values. Genotypes were generated under a mutation rate *μ* = 10^−8^ using a simulated ARG covering a 10-kb gene region for 20,000 individuals under a GBR demographic model. Under the null, phenotypes were simulated as standard normal. Under the alternative, 10% of variants were set as causal with effect sizes drawn from 𝒩(0, [*f*(1 − *f*)]^−1^),where *f* is the minor allele frequency, and noise was added to achieve *h*^2^ = 0.02.

Supplementary Figure S4 shows the distribution of test statistics under varying levels of missingness (2%, 5%, and 10%). Under the null, missingness had minimal effect on the distribution of *T*, confirming that type I error is well controlled. Under the alternative, randomly missing values introduced a downward bias in the test statistic, while selectively missing extreme values led to inflation. Since this inflation occurs under the alternative, it does not result in false positives but reflects uncalibrated *p*-values; in contrast, deflation from random missingness may lead to a reduction in power.

Finally, we note that under the alternative, the phenotype vector becomes correlated with the GRM *R*_11_ even when entries are missing at random, as the spectrum of the observed GRM depends on the non-missing subset of individuals. As a result, there is no general guarantee on the direction or magnitude of the bias relative to the fully observed case, an issue that affects all methods based on covariance structure testing, including ours. We refer to [4] for further theoretical details.

These results confirm that imputing missing values as 0 preserves calibration under the null and is overall more conservative than excluding individuals, though the impact on power depends on missingness patterns.

### 1.3 Effect of mutation sampling rate on association power

ARG-RHE performs association testing using mutations resampled from the inferred ARG. As discussed in [5], this provides a Monte Carlo approximation to the full ARG-GRM. Here, we assess how the mutation resampling rate *μ* affects statistical power and computational cost.

Using a fixed sample size of 100,000 individuals and *h*^2^ = 0.01, we simulated phenotypes and causal variants as in Figure 3 of the main text. We varied *μ* from 10^−8^ to 5 × 10^−6^ and computed association power in 10-kb gene regions. As shown in Supplementary Figure S2(a), power increased with *μ*, but plateaued near *μ* = 10^−6^.

Supplementary Figure S2(b) shows the corresponding CPU time and memory usage. ARG-matrix multiplication typically scales sublinearly due to redundancy among nearby mutations. However, at high mutation rates, repeated hits to the same edges increase marginal costs, leading to a threshold effect around *μ* ≳ 2 × 10^−6^. To balance power and efficiency, we chose *μ* = 10^−6^ for our UK Biobank analyses.

## 2 Additional details on ARG-matrix multiplication

This section provides some background on ARG representation and traversal, and further algorithmic details on ARG-matrix multiplication and its application in ARG-based LMM analyses.

### 2.1 Background: ARG representation and traversal

#### ARG representation

The ancestral recombination graph (ARG) is a graph (*N, E*) in which nodes *N* represent genetic sequences from sampled individuals and their ancestors, and edges *E* encode the inheritance of genomic segments through time and along the genome. ARG nodes and edges also carry meta-information related to time and genomic coordinates: each node *n* ∈ *N* is annotated with an age (e.g., in generations), and each edge *e* ∈ *E* is associated with a genomic interval, denoting the region transmitted from a parent to a child. Nodes with no descendants, often referred to as leaves, typically represent sampled individuals, while nodes with no ancestors, referred to as roots, typically represent the most recent common ancestors for all samples within a region of the genome. Mutations are assigned to edges of the ARG at specified positions and times at which they first arose in the sample. The descendants of a mutation are the leaf nodes reachable through directed paths from the mutation-bearing edge. Conversely, the genotype of a sample can be assembled by collecting all mutations lying on edges ancestral to its corresponding leaf node.

ARGs have been extensively studied [6–21], with significant advances in their computational handling driven by the development of genomic simulators [10, 11, 13–21]. A key property of ARGs is that they reduce to a marginal tree when only the nodes and edges overlapping a given genomic position are retained. These marginal trees, however, are only implicitly represented, as many nodes and edges are shared across neighboring positions and thus appear in multiple marginal trees. Although some simulators, particularly those processing the genome from left to right, use marginal-tree-based representations, others adopt more memory-efficient graph-based structures that explicitly map ARG nodes and edges to concrete elements of a data structure, while balancing other computational trade-offs [11, 15, 17, 19–21]. For example, COSI2 [17] tracks only the subset of ARG nodes that are active at a given time to reduce memory usage. However, in order to output mutations as they are sampled, it also maintains the set of descendants for each active region, which results in substantial memory costs for large sample sizes. ARGON [19] and msprime [20] independently introduced graph-based ARG representations and traversal strategies that support efficient mutation placement and analysis without the need to store large descendant sets in memory. In this work, as done in ARGON and elsewhere [5, 19, 22], we refer to this graph-based data structure as an ARG. A similar representation is referred to as a “tree sequence” by the developers of msprime and tskit [20, 21, 23, 24].

#### ARG traversal

Efficient traversal algorithms allow ARG-based summaries to be computed while holding only a small amount of state in memory. To illustrate the underlying ideas, we summarise the traversal used in ARGON [19] (Algorithm S1) to generate mutations, compute the expected allele frequency spectrum (AFS), and identify pairwise identical-by-descent (IBD) segments. In ARGON, individual ARG nodes are represented as lists of “blocks”, contiguous genomic intervals that point to intervals in other nodes, acting as edges between them. For ease of exposition, we refer to these blocks simply as “nodes” in the pseudocode of Algorithm S1, but use “node block” in this text to emphasize that they represent intervals within ARG nodes. The algorithm combines a top-down discovery pass with a bottom-up processing pass. For each ARG root (a node block with no parents), processed from left to right along the genome, a top-down traversal is first performed to mark all reachable descendant node blocks. These are then enqueued in a priority queue, sorted by their time, and processed from recent to deep time. At each block, the set of descendant leaves is memoized as the union of the descendant sets of its children (Line 16), which have been previously computed, avoiding redundant computation. These sets are used to (i) generate mutations on incident edges, (ii) accumulate branch area into the appropriate AFS bin, and (iii) emit IBD segments defined by pairs of lineages that coalesce at the current block. Once a child block has been used by all of its parents, its descendant list is discarded, limiting peak memory usage.

##### Algorithm S1

ARGON [19] ARG traversal to compute mutations, expected AFS, and pairwise IBD^1^

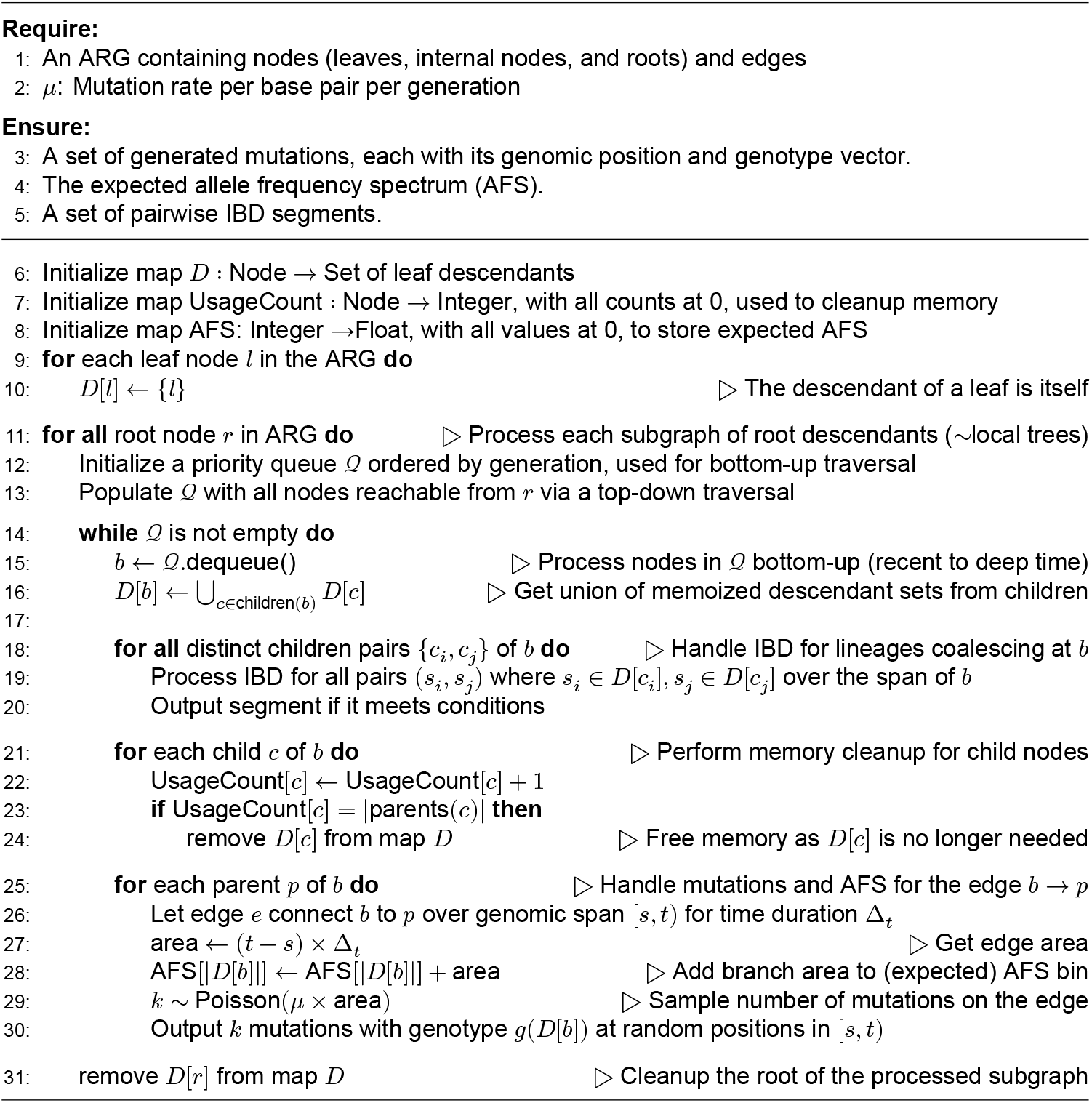

This type of graph traversal has been leveraged for efficient computation in several efficient ARG-based frameworks. A related approach was independently developed in msprime [20]. It was later extended within tskit to compute various statistics, including *F* -statistics [25] and IBD sharing [26]. In `arg-needle-lib`[5], where ARG-RHE is implemented, ARG traversals are used for computations relevant to complex trait analysis, including the construction of ARG-GRMs for LMM analyses. These GRMs are computed by accumulating the volume of edges with specific properties, analogous to how edge volumes are summed in Algorithm S1 to compute the expected AFS. A subtle distinction between the traversal algorithm in Algorithm S1 and that used in `arg-needle-lib` stems from the way recombination is handled in ARGON, where node blocks have, by construction, a fixed set of children and a fixed descendant set, a property referred to as a “bricked” genealogy in [27]. This slightly increases the number of edges but enables constant-time access to a node block’s children. In contrast, `arg-needle-lib` relaxes this constraint, allowing a node span to have different descendant sets along its interval. This can reduce the number of nodes, at the cost of slightly increased computation to determine the correct descendant set at each position, using an interval tree (also used in [17, 20]). As a result, the traversal logic is adapted to track the nearest genomic location at which the children of traversed nodes may change (an “expiry position”), maintaining these positions in a separate priority queue used to determine when memoized results should be discarded or recomputed. This logic is used to implement several types of traversal functions with slightly different properties, available at https://github.com/palamaraLab/ `arg-needle-lib`, including `visit_branches, visit_clades`, and `visit_mutations.` The latter, whose pseudocode is shown in Algorithm S2, is used to efficiently traverse the ARG and compute quantities associated with mutations on ARG edges, and is leveraged for left-side ARG-matrix multiplication, described below.

##### Algorithm S2

arg-needle-lib [5] traversal for mutation-based statistics (visit_mutations)^2^

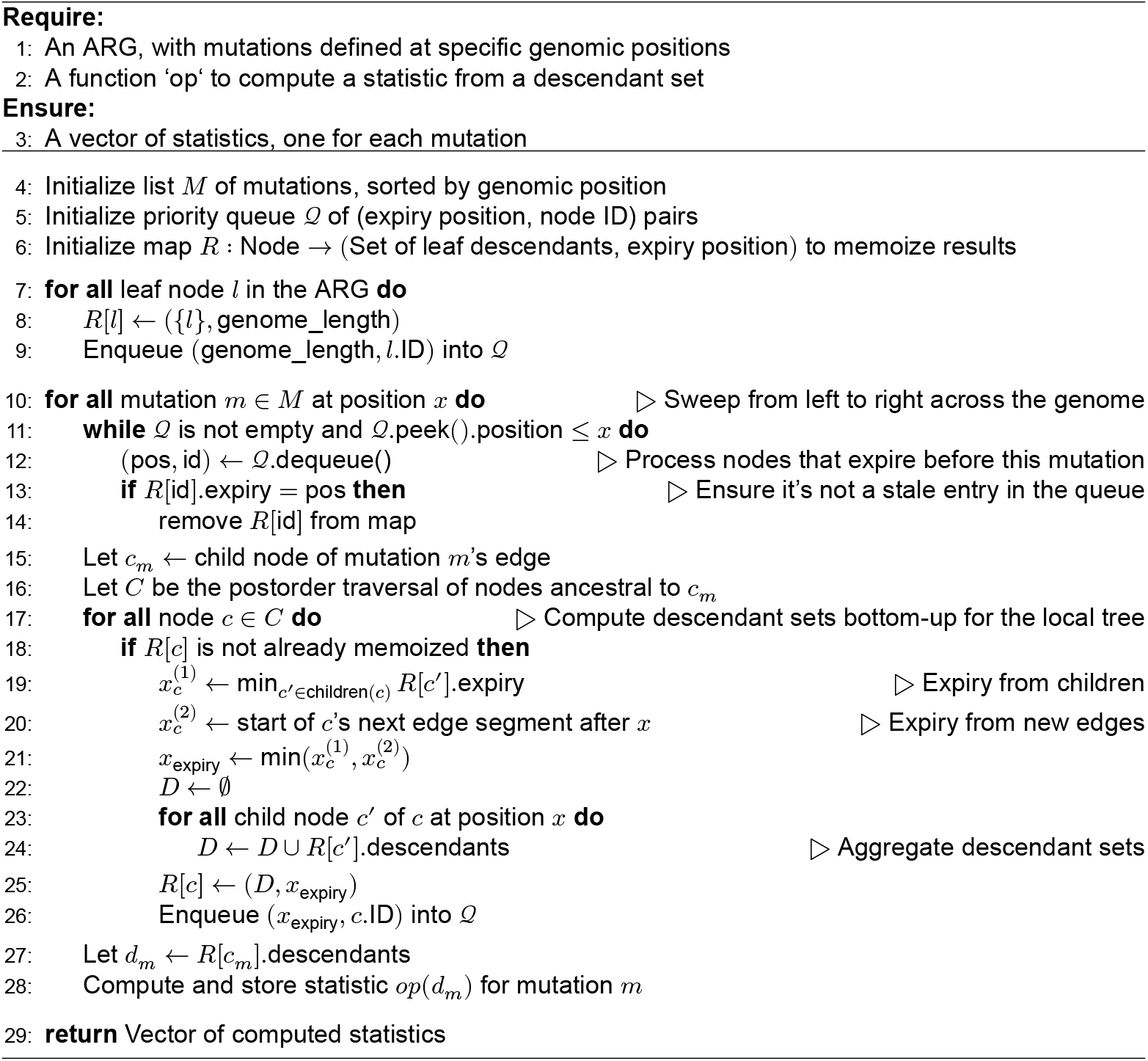

### 2.2 Left-side multiplication: *V* = *UX*

Building on these approaches, we now describe ARG-matrix multiplication, where the goal is to compute *V* = *UX*, with *X* denoting the *n* × *p* genotype matrix and *U* an input matrix multiplied on the left. An early version of this in-ARG multiplication approach was described in [28].

Let *x* ∈ 0, 1, 2^*n*^ denote the vector indicating which individuals carry a particular mutation; then a column of *V* is given by *v* = *U x*. As with other computations described above, the hierarchical structure of the ARG implies that the descendants of a segment represented by an ARG edge are the union of the descendants of its child edges. Accordingly, the carrier vector *x* for an edge can be defined recursively as *x* = ∑_*c*_ *x*_*c*_, where *x*_*c*_ is the carrier vector for child edge *c*. This allows *v* = *U x* to be computed as *v* = ∑_*c*_ *U x*_*c*_, with partial products *U x*_*c*_ memoized to enable efficient traversal and accumulation.

Algorithm S3 implements this bottom-up traversal. It follows the logic of Algorithm S2, where the op function is configured to perform matrix–vector products. The algorithm ensures that all required descendant information is available for each mutation before aggregation proceeds upward. To handle variation in descendant sets along a node’s genomic interval, it tracks an expiry position for each cached result. These expiry positions indicate genomic locations where the cache becomes invalid, either due to a recombination event in a child or the appearance of a new child edge. The traversal purges expired results and updates them accordingly.

#### Algorithm S3

Left-side ARG-matrix multiplication (*V* = *UX*)

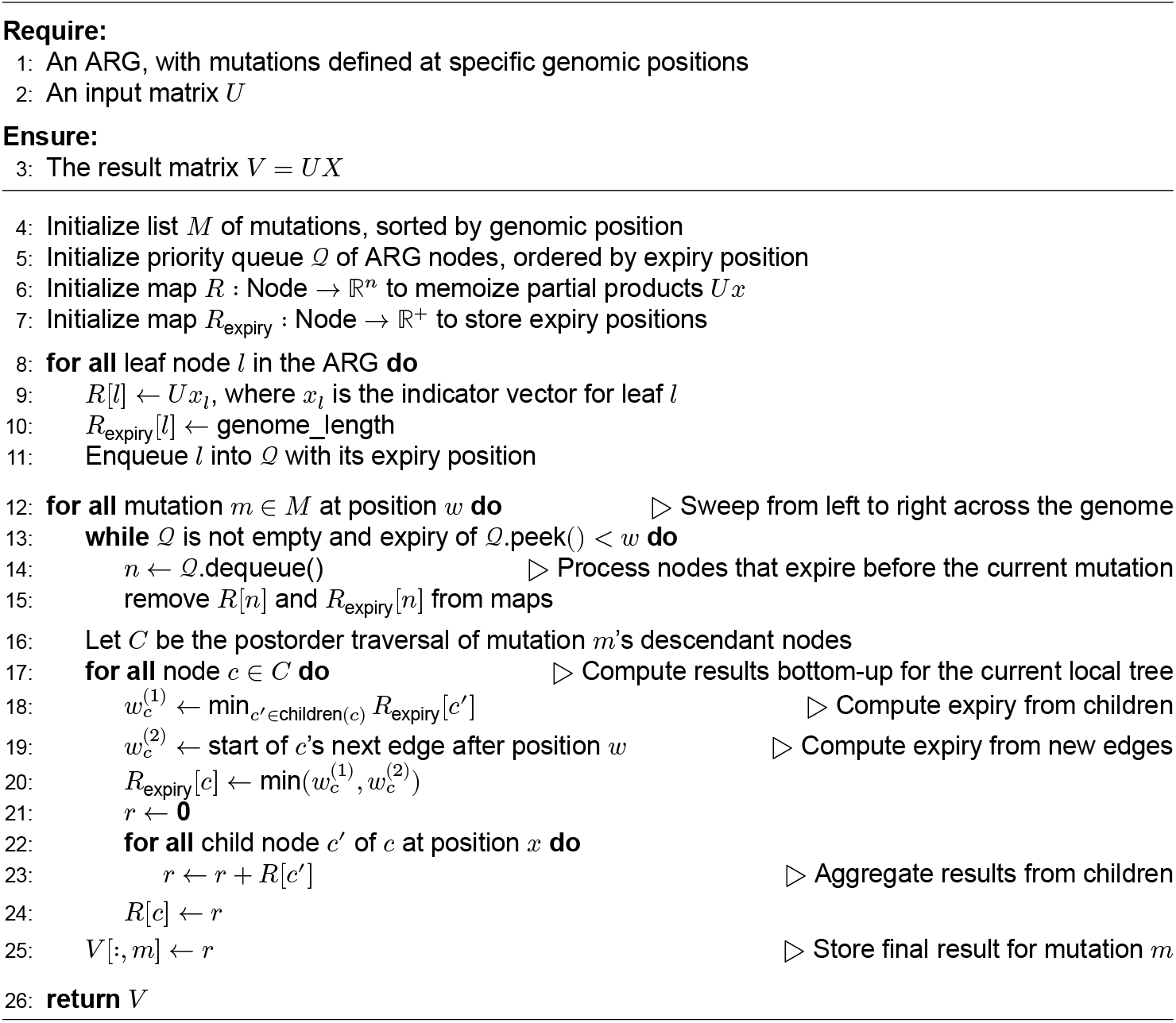

### 2.3 Right-side multiplication: *V* = *XU*

We now consider computing *V* = *XU*, where *U* is an input matrix multiplied on the right. This operation aggregates information across mutations for each node and requires a top-down traversal of the ARG from roots to leaves. Let *x* denote the genotype vector for a given node; then *v* = *xU* . As before, genotype vectors can be expressed recursively in terms of ancestral segments. For nodes with multiple parent edges due to recombination, the overall genotype is a sum of contributions from each parent segment. This gives *v* = ∑_*p*_ *x*_*p*_*U*, where *x*_*p*_ is the genotype vector corresponding to parent edge *p*.

Algorithm S4 implements this top-down traversal. Unlike the left-side case, expiry positions are now precomputed in a prior pass, since the direction of inheritance is reversed. Genotype information is cached for each contiguous genomic block bounded by recombination events and is propagated from ancestors to descendants. Recombination is again tracked explicitly to avoid prematurely merging results across incompatible genomic regions.

#### Algorithm S4

Right-side ARG-matrix multiplication (*V* = *XU*)

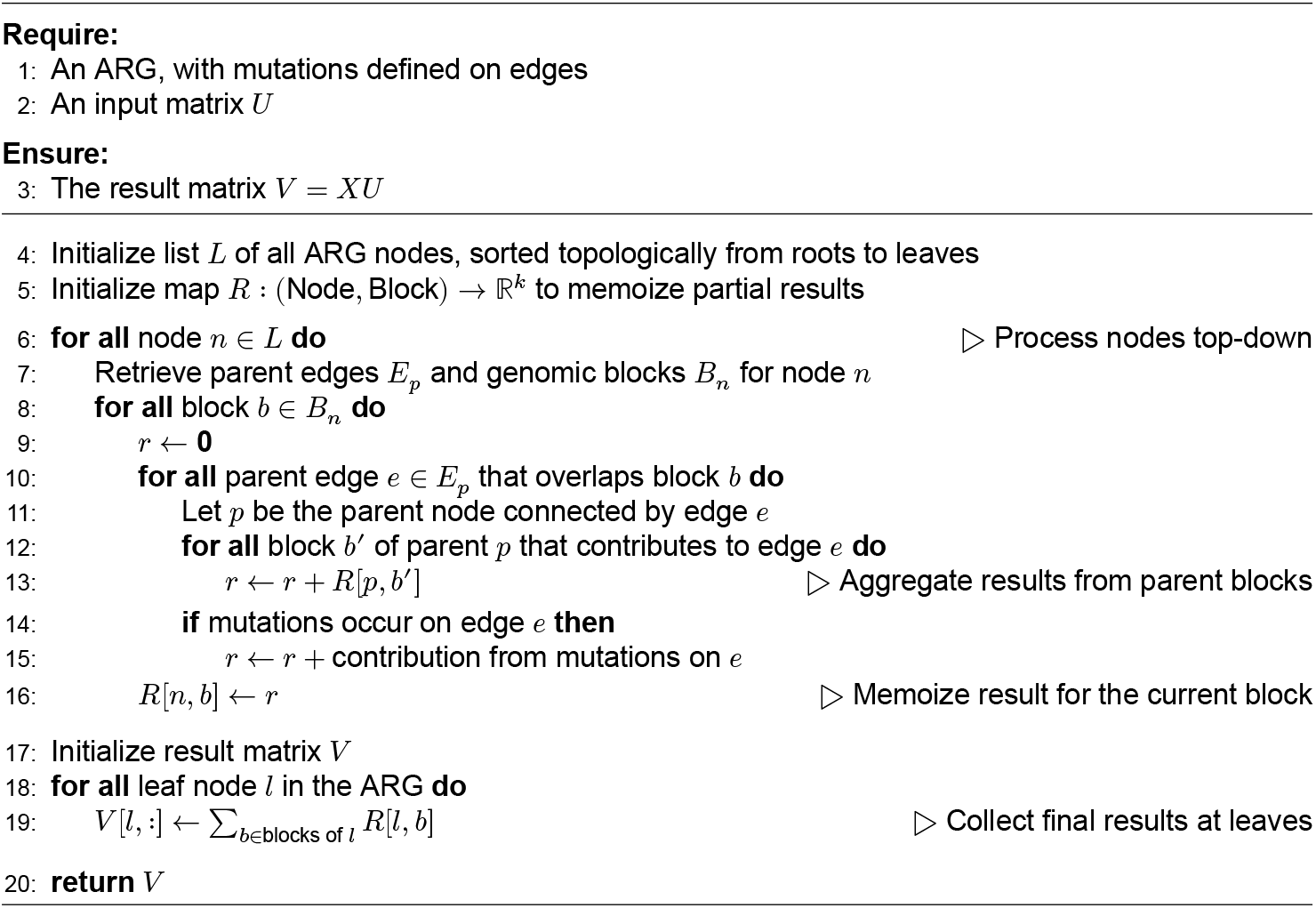

### 2.4 BLUP and LOCO calculations for ARG-based prediction and association

In addition to estimating heritability and performing variance component association testing, ARG-matrix multiplication can accelerate other ARG-based LMM analyses described in [5], including polygenic prediction and association testing. This includes the efficient computation of best linear unbiased predictors (BLUPs), which are also used to generate residualized phenotypes for association. We describe this algorithm below, following the ARG-based conjugate gradient strategy also described in [28].

Several LMM-based GWAS methods [29–33] rely on a two-step procedure. In step 1, phenotypes *Y* are regressed on covariates *W* and a set of genetic variants 𝒢_1_ to correct for population structure and polygenic background. In step 2, a separate set of variants 𝒢_2_ is tested for association with the residual phenotype, typically using a leave-one-chromosome-out (LOCO) scheme: for each chromosome *c*,variants on *c* are tested against 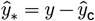 where 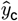 is the prediction from step 1 excluding variants on chromosome *c* [34].

We follow the algorithmic strategy used in BOLT-LMM-Inf, where 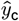 is computed as the best linear unbiased predictor (BLUP) [29], now leveraging the ARG to enable scalable BLUP calculation using ARG-based GRMs (see also [28]). The prediction 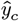 is proportional to 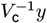, with 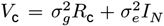 the LOCO covariance matrix. Rather than inverting *V*_c_, we solve the system *V*_c_*x* = *Y* using conjugate gradients (CG) [28, 29, 35], an iterative method that only requires matrix–vector products. Each iteration requires computing 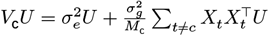, where *X*_t_ is the genotype matrix for chromosome *t* and *M*_c_ is the total number of variants on all chromosomes except *c*. ARG-matrix multiplication enables efficient evaluation of these products without explicitly forming the GRM, using Algorithms S3 and S4 on each chromosome and accumulating the results on the fly, as described in Algorithm S5. Empirically, the number of CG iterations grows roughly with 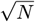 [29]. The primary bottleneck in this calculation is the repeated evaluation of residual vectors 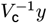 . A comparison between a NumPy-based implementation of Algorithm S5 and one using ARG-matrix multiplication is shown in Figure 1e.

#### Algorithm S5

Calculate phenotype residuals 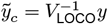 using conjugate gradients

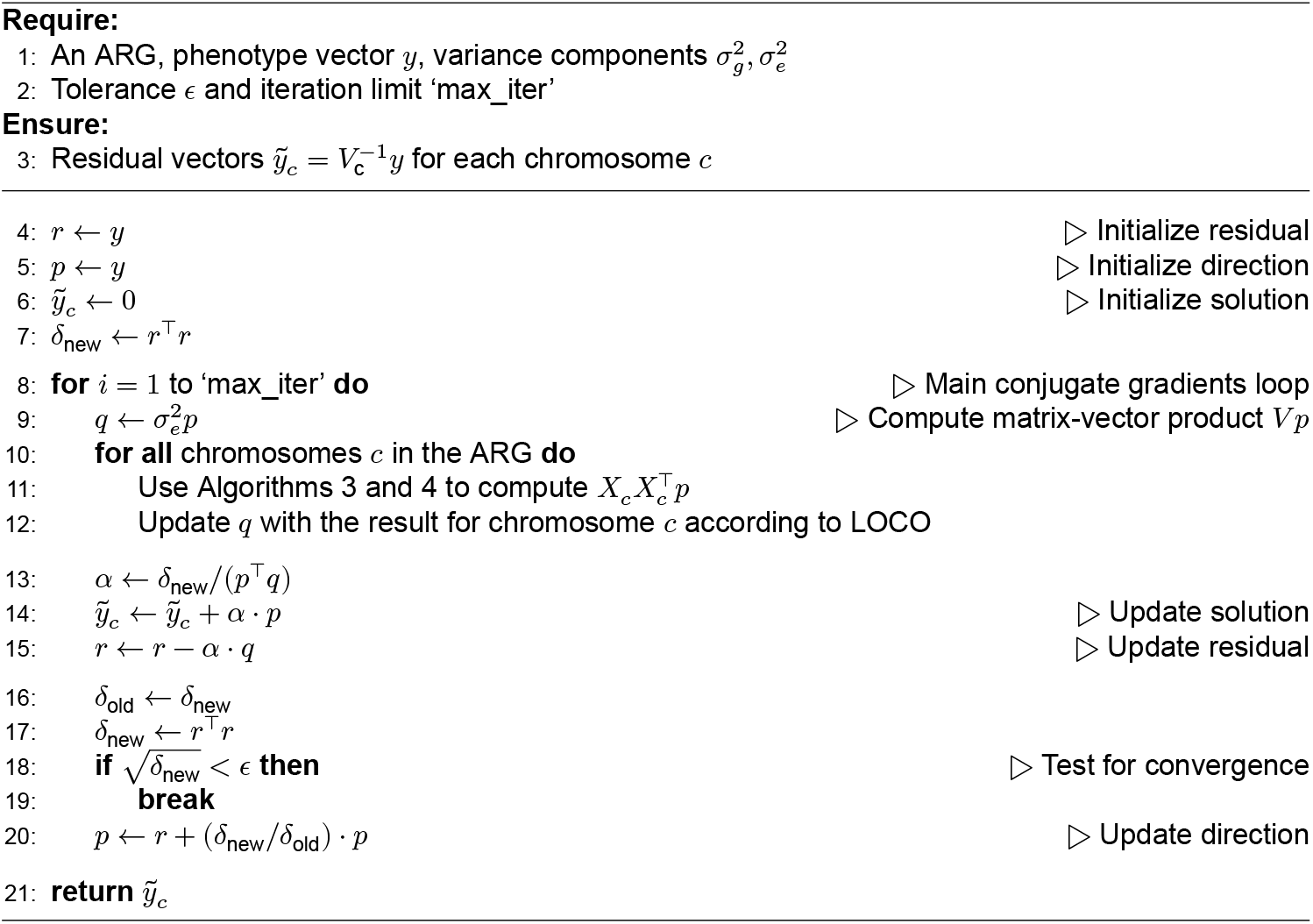

We note that, although not shown for simplicity, lines 13–17 can be parallelized to simultaneously analyze multiple chromosomes and phenotypes. The resulting LOCO-residual phenotype vector 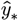 can then be tested for association against each variant *g* ∈ 𝒢_2_ using the GRAMMAR-GAMMA approximation to the score test statistic 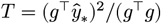 [29, 36]. Additional details may be found in [28].

## 3 Additional experimental details

This section provides additional details related to our UK Biobank analyses.

### 3.1 Controlling for population structure in UK Biobank analyses

To assess the effectiveness of controlling for population stratification using 20 principal components (PCs), we applied ARG-RHE to 337,464 white British individuals in the UK Biobank. We simulated 50 heritable phenotypes (*h*^2^ = 0.5), each generated by selecting 50% of all imputed variants on the odd-numbered chromosomes as causal. Effect sizes were drawn from a normal distribution with variance proportional to 1/(*f*[1 − *f* ]), where *f* denotes the minor allele frequency, and independent Gaussian noise was added to achieve the target heritability. Association testing was then performed using inferred ARGs covering gene regions on the even-numbered (non-causal) chromosomes, after residualizing phenotypes against the top 20 PCs computed in [37].

As shown in Supplementary Figure S3, the quantile-quantile (Q-Q) plot of *p*-values from non-causal regions closely follows the expected uniform distribution, confirming effective population structure correction.

### 3.2 Assessing tissue relevance using gene expression data

To assess the biological relevance of gene-based associations, we examined whether identified genes are preferentially expressed in a relevant tissue. We focused on 27 blood-related phenotypes (Supplementary Table 2), using whole blood as the reference tissue.

We used the `GENE2FUNC` module in FUMA [38] to annotate tested genes with normalized expression values across 54 tissues from GTEx v8. For each gene, we selected the five tissues with the highest expression levels, considering only those with a normalized TPM *z*-score above 1. A gene was labeled “relevant” if whole blood was among its top five tissues.

We then computed the proportion of relevant genes identified by ARG-RHE, HRC-RHE, and ACAT-V, averaging across chromosomes to estimate standard errors. As a baseline, we repeated the same procedure using a random selection of genes and phenotypes.

As shown in Supplementary Figure S9, ARG-RHE yielded a relevance rate of 23.6%, compared to 19.6% for the baseline (*t*-test, *p* = 1.56 × 10^−4^). HRC-RHE and ACAT-V yielded similar rates of 24.6% and 24.4%, also significantly enriched relative to the baseline (*t*-tests, *p* = 3.95×10^−5^ and *p* = 4.03×10^−6^, respectively), but not significantly different from ARG-RHE (*p* > 0.05).

## Footnotes

1 Pseudo-code obtained using Gemini 2.5 Pro based on the ARGON source code at https://github.com/pierpal/ARGON/.

2 Pseudo-code obtained using Gemini 2.5 Pro based on the source code at https://github.com/PalamaraLab/arg-needle-lib/.

